# The response network of HSP70 defines vulnerabilities in cancer cells with the inhibited proteasome

**DOI:** 10.1101/2021.11.16.468807

**Authors:** Magdalena Oroń, Marcin Grochowski, Akanksha Jaiswar, Justyna Legierska, Kamil Jastrzębski, Magdalena Nowak-Niezgoda, Małgorzata Kołos, Wojciech Kaźmierczak, Tomasz Olesiński, Małgorzata Lenarcik, Magdalena Cybulska, Michał Mikuła, Alicja Żylicz, Marta Miączyńska, Katherina Zettl, Jacek R. Wiśniewski, Dawid Walerych

**Affiliations:** Mossakowski Medical Research Institute PAS, Warsaw, Poland; International Institute of Molecular and Cell Biology, Warsaw, Poland; Central Clinical Hospital of Ministry of Interior and Administration, Warsaw, Poland; National Center of Oncology, Warsaw, Poland; Max Planck Institute of Biochemistry, Martinsried, Germany

**Keywords:** HSP70, proteasome, carfilzomib, proteomics, cancer

## Abstract

Human neoplasias are often addicted to the proteasome machinery. However, cancers have evolved efficient response mechanisms to overcome proteasome inhibition with bortezomib and carfilzomib - drugs approved for multiple myeloma treatment. To understand these responses we investigated proteome changes upon the proteasome inhibition with carfilzomib - in multiple myeloma, normal fibroblasts, and cancers of lung, colon, and pancreas. A pathway-oriented siRNA screen based on the proteomics results showed that molecular chaperones, autophagy- and endocytosis-related proteins are cancer-specific vulnerabilities combined with carfilzomib. Targeting of HSPA1A/B (HSP70 family chaperones) most specifically sensitized cancer cells and patient-derived organoids to the proteasome inhibition. A high level of HSPA1A/B mRNA correlated with a low proteasome activity in cancer patient tissues and is a risk factor in cancer patients with a low proteasome expression. Mechanistically, HSPA1A/B governed autophagy, unfolded protein response, endocytic trafficking, and chaperoned the proteasome machinery, suppressing the effect of the proteasome inhibition, but did not control the NRF1/2-driven proteasome subunit transcriptional bounce-back. Consequently, downregulation of NRF1 most specifically decreased the viability of cancer cells with the inhibited proteasome and HSP70.

## Introduction

Proteasome machinery is the central player in the controlled cellular protein degradation in eukaryotic cells ^1^. It consists of the 20S catalytic core capable of degrading a significant proportion of cellular proteins independently of ubiquitination ^2^, supplemented by 19S regulatory caps providing a substrate polyubiquitination dependence ^3^. Specific core subunits are replaced by alternative proteins in immunoproteasomes – responsible for peptide processing in antigen presentation ^4^. The catalytic core of all these proteasome forms possesses three main proteolytic activities in the 26S proteasome termed: chymotrypsin-like, trypsin-like and caspase-like. The first of them is rate-limiting ^5^, and thus its targeting with inhibitors affects major proteasome functions in cells ^6^.

Neoplastic cells are addicted to the activity of the proteasome machinery, which enables degradation of tumor suppressors and allows to sustain protein homeostasis in transformed cells ^7,8^. Therapeutic proteasome inhibitors, pioneered by bortezomib, have been successfully utilized in multiple myeloma treatment ^6^. However, they were found relatively ineffective in solid tumors in clinical trials ^9^, while multiple myeloma patients have frequently developed resistance to the bortezomib treatment ^10^. Thus, second-generation inhibitors were developed, including carfilzomib, approved for a relapsed/refractory multiple myeloma treatment ^11^. The drug is more specific than bortezomib, effective towards the 20S, 26S, and immunoproteasome by irreversibly inhibiting the chymotrypsin-like proteasome activity, and at higher concentrations – the caspase- and trypsin-like activities ^12–14^. Nevertheless, the introduction of carfilzomib has not resolved the problem of cancer’s resistance to proteasome inhibition, suggesting the existence of effective cancer-specific compensatory responses ^15^.

Several mechanisms have been put forward as the compensatory responses to proteasome inhibition in neoplasias. Expression of 26S proteasome subunits is increased during a bounce-back response to proteasome inhibitors, mediated mainly by NRF1 (NFE2L1) ^16^ and NRF2 (NFE2L2) optionally in conjunction with mutant p53 ^17^. Components of the autophagy-lysosome protein degradation system are upregulated on proteasome inhibition ^18,19^. Concomitant inhibition of proteasome and autophagy pathways was beneficial in treating models of several cancer types, including breast, pancreatic and hepatocellular carcinoma ^20–22^. Proteins accumulating on the proteasome inhibition induce an unfolded protein response (UPR), which leads to context-dependent cell death or survival, which can be both affected by drugs targeting UPR in neoplasias ^23–26^. Accumulation of misfolded proteins leads also to the induction of molecular chaperone proteins, including members of HSP70, HSP90 families of chaperones, and DNAJ and BAG families of co-chaperones ^27–30^. Targeting these proteins was considered to increase the effects of proteasome inhibitors in neoplasias ^31–33^. However, the extent and mechanisms of chaperone response to the proteasome inhibitors have not been elucidated.

In this study, we employed a global proteomics approach to compare in an unbiased manner the contributions of different compensatory mechanisms upon the proteasome inhibition with carfilzomib in cells of multiple myeloma, normal fibroblasts, and cancers of lung, colon, and pancreas. The cancer types were chosen based on the highest numbers of deaths they are predicted to cause in the upcoming decade among neoplasias of men and women altogether, in the US and the EU countries ^34,35^. We defined which elements of the response center around the main HSP70 family stress-inducible molecular chaperones HSPA1A and HSPA1B - the most consequent proteasome inhibition responders in proteomes of all the studied cell types (often addressed jointly as HSPA1A/B thanks to nearly identical protein sequences ^36^). This led to the identification of a network of processes co-activated with HSP70, controlled by HSP70, and independent of HSP70 in response to proteasome inhibition. We found that HSPA1A/B induction in cancer cells supports autophagy, UPR, endosomal transport, and directly - the 26S proteasome activity. However, HSPA1A/B did not support NRF1/2-dependent proteasome bounce-back, opening a possibility to target this pathway in parallel to HSP70.

## Results

### Proteomics reveals similarities and differences in proteasome inhibition response in cancer, multiple myeloma, and normal fibroblasts

In an attempt to find a therapeutic window to specifically kill cancer cells using proteasome inhibition, we first compared their response to carfilzomib with multiple myeloma and normal cells. We tested viability in pairs of cell lines from three cancer types – lung, colon, and pancreatic – against pairs of multiple myeloma cell lines and primary human fibroblasts, 24h post-treatment with increasing concentrations of carfilzomib (Fig. 1A). Multiple myeloma cell lines were the most sensitive to the proteasome inhibition, with a calculated IC50 at 8-12 nM of carfilzomib, while in the cancer cell lines IC50 was in a range of 66-326 nM and for normal fibroblasts – 424-512 nM of carfilzomib (Fig.1A, Table S1). Hence, cancer cells remained more sensitive to carfilzomib than normal cells, while multiple myeloma cells were severalfold more sensitive than either of them. To investigate if this phenomenon was mirrored by changes in the proteasome activity, we determined carfilzomib’s IC50s of the proteasome chymotrypsin-like activity in the same set of cell lines. In this experiment, the normal fibroblast cells were the most sensitive to carfilzomib, while the cancer cells were on average the least sensitive (Fig. 1B, Table S1). As a result cancer cells and normal fibroblasts on average retained nearly 100% viability at 50% of the proteasome activity upon carfilzomib treatment for 24h, which was significantly higher than the multiple myeloma cells’ viability (Fig. 1C). However, cancer cells had a significantly lower chymotrypsin-like proteasome activity at the 50% of their viability than the multiple myeloma cells, while it was significantly higher than the normal fibroblasts (Fig. 1D). This suggested that cancer cells possess response mechanisms that allow them to survive low proteasome activity similarly to normal cells and unlike multiple myeloma, while they additionally retain more proteasome activity than the normal cells under the carfilzomib treatment.

**Figure 1.**
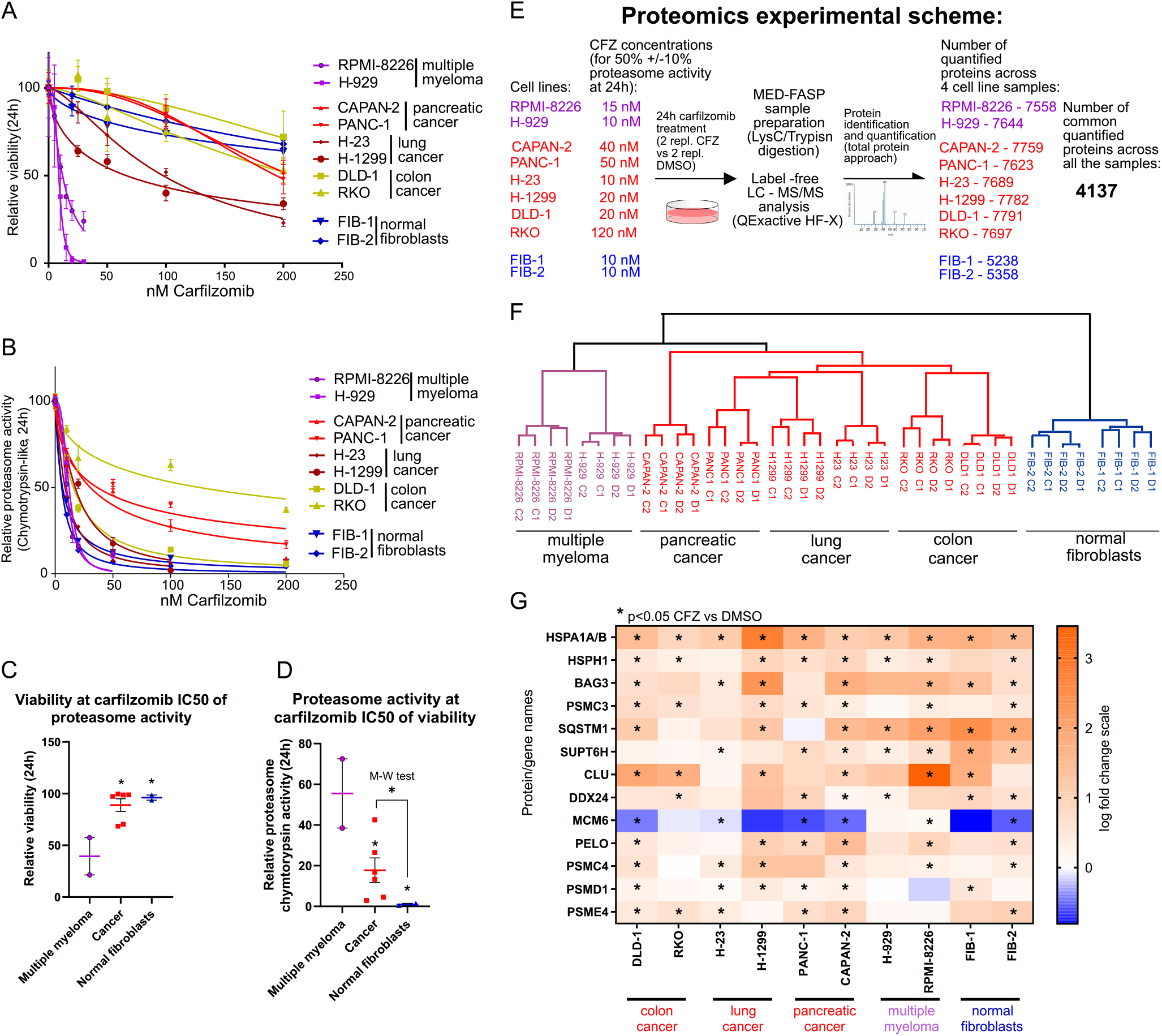
Differences and similarities of cancer, multiple myeloma and normal fibroblast cells response to the proteasome inhibition. **A,** Indicated cell lines were treated with increasing concentrations of Carfilzomib for 24h, their viability was measured by the ATP-lite assay (Methods) and normalized results plotted against the used Carfilzomib concentrations (means of n=3, with SD), with nonlinear least-squares regression curves fitted to the data points. **B,** Indicated cell lines were treated with increasing concentrations of Carfilzomib for 24h, and their chymotrypsin-like proteasome activity was measured (Methods) and normalized results plotted against the used Carfilzomib concentrations (means of n=3, with SD), with nonlinear least-squares regression curves fitted to the data points. **C,** Differences in viability of indicated cell line groups calculated at Carfilzomib IC50 of the proteasome activity from B (details in Table S1). Means with SEM, one-way ANOVA with Dunnett’s correction, * p<0.05. **D,** Differences in chymotrypsin-like proteasome activity of indicated cell line groups calculated at Carfilzomib IC50 of the viability form A (details in Table S1). Means with SEM, one-way ANOVA with Dunnett’s correction, * p<0.05. For cancer vs. multiple myeloma cells Mann-Whitney test was additionally performed - * p<0.05. **E,** Scheme of proteomics experiment performed using samples from shown cell lines treated with indicated carfilzomib concentrations (for 50% ±10% proteasome activity) and listed numbers of quantified proteins. **F,** Hierarchical clustering (Euclidean distance) of proteomics results for 4137 proteins quantified in each of 40 proteomes from 10 cell lines (proteins not quantified in at least one sample removed). **G,** Heat map of log fold changes in differential protein level analysis between CFZ (carfilzomib treated) and DMSO (control) in proteomes of indicated cell lines. Proteins with t-test p-value<0.05 significant result are marked with *. Only proteins with 6 or more significant results are shown. Full result available in Table S3.

To understand molecular mechanisms underlying the specific features of cancer, multiple myeloma, and normal cell responses to the proteasome inhibition, we performed a global proteomics analysis in lysates of the ten cell lines used in the experiments described above (Fig. 1E). The cells were treated with concentrations of carfilzomib which inhibited the chymotrypsin-like proteasome activity to 50% (+/- 10%) at 24h post-treatment (Fig. 1E; carfilzomib concentrations were based on the calculated IC50s of the proteasome activity from the Table S1, adjusted to the experimentally measured proteasome activity values). A label-free mass spectrometry analysis and a global protein approach with the sensitivity of over ten thousand identified proteins in a sample ^37,38^ allowed for identification and quantification, across all four samples analyzed for each cell line, of more than seven thousand proteins in multiple myeloma and cancer cells and more than five thousand proteins in normal fibroblasts, (Fig. 1E, Fig. S1A, Table S2). 4137 proteins, identified and quantified in each of the 40 samples analyzed overall, were used for hierarchical clustering analysis with Euclidean distance. This analysis revealed that the proteomes of the normal fibroblasts, cancer, and multiple myeloma cells cluster separately (Fig. 1F). Differential analysis of protein level changes in carfilzomib-treated samples versus the DMSO-treated controls in each cell line (Table S2) showed only one hit significantly changing level (p<0.05) across all the cell lines – the main stress-inducible HSP70 family proteins, unified in the proteomics result due to a very high sequence similarity - HSPA1A/B (Fig. 1G, Fig. S1B). No specific proteins differed significantly in the six or five cancer cell lines *versus* the two normal fibroblasts and the two multiple myeloma cell lines (Fig. 1G, Table S3). Most proteins significantly changing levels across multiple analyzed cell lines were either upregulated molecular chaperones or upregulated proteasome subunits (Fig. 1G). This suggested that the overall response to the proteasome inhibition is affecting similar proteins in different cell types, while the result of the hierarchical clustering of the studied proteomes indicated that broader, systems-biology analysis of the proteomics data is required to hunt for cancer-specific proteasome compensators.

### Molecular chaperone, autophagy, and endocytosis-related proteins are vulnerabilities in the carfilzomib-treated cancer cells

To confront systemic responses to proteasome inhibition in neoplastic and normal cell line groups we performed a comparative pathway analysis between changes in their proteomes. The proteins significantly changing levels (p<0.05) upon carfilzomib treatment in cancer, multiple myeloma and normal fibroblast cell lines were each fused into signatures, filtered for duplicates, and each signature was analyzed separately for enriched molecular pathways with association FDR<0.05, followed by the resulting overlap aimed to find common and specific pathways (Fig. 2A, Table S4). The ClueGO tool analysis revealed cancer-specific pathways which represented cellular functions such as translation, mitochondrial metabolism, autophagy, UPR, MAPK signaling, or endocytosis, as well as pathways common to cancer, multiple myeloma, and/or normal cells (Fig. 2A). Similar pathways and functional protein groups were found by the Ingenuity Pathways Analysis (Fig. S2A). The pathways common to all three cell lines types were strongly enriched in the proteasome subunit proteins (Table S4), reflecting the bounce-back response to the proteasome inhibition (Fig. 2A). Genes encoding proteins from functional groups significantly up or downregulated in cancer cells found in pathways specific to cancer and common to cancer and multiple myeloma or normal fibroblasts, indicated in Figure 2A, were used as targets in a cancer vulnerability siRNA mini-screen. The choice of particular genes/proteins for the mini-screen was based on the presence in the cancer cells’ signature, level change compared to multiple myeloma or normal fibroblasts, and the availability of candidate therapeutics for potential repositioning in supplementing the proteasome inhibitor-based therapy (listed in Table S4).

**Figure 2.**
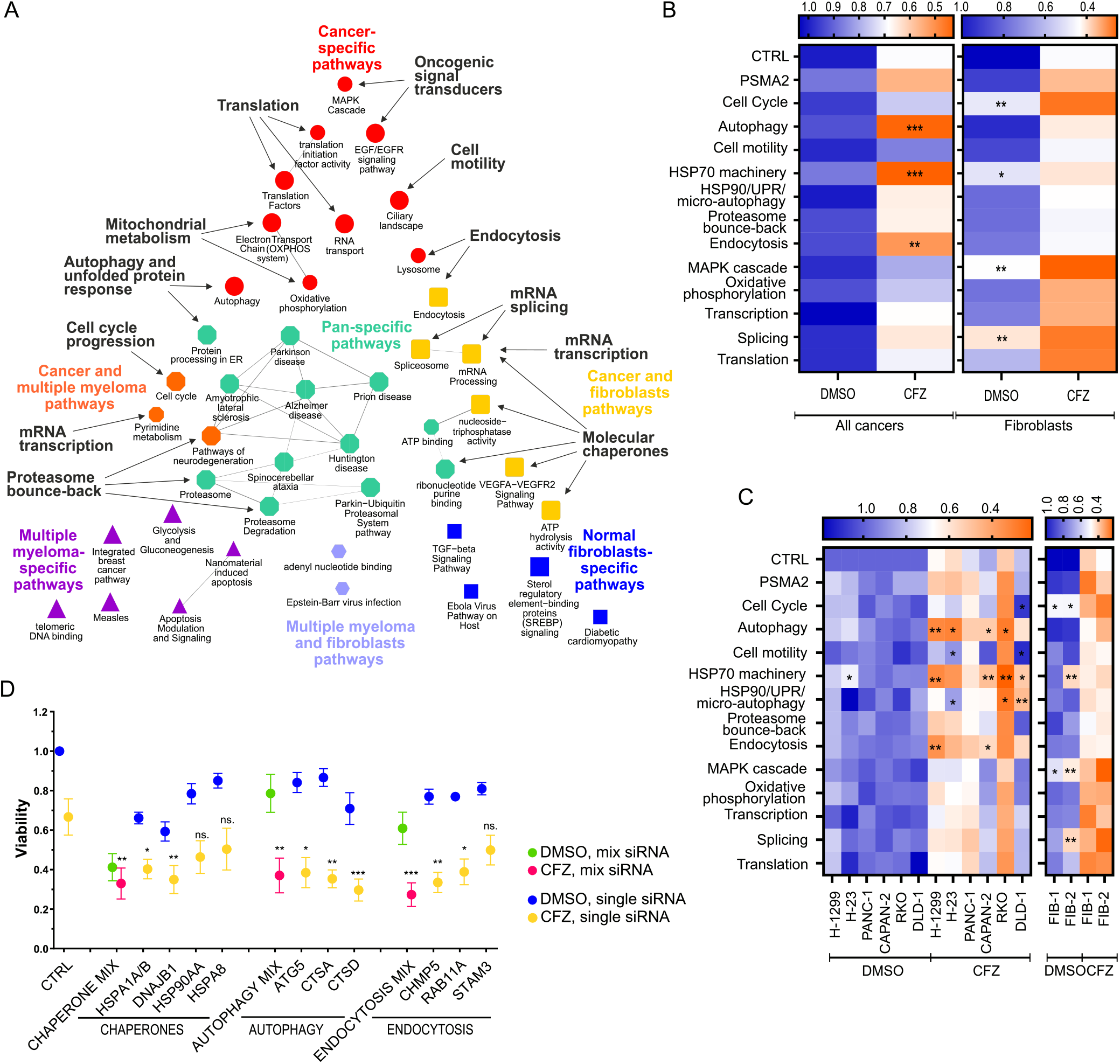
Molecular pathways involving molecular chaperones, autophagy, and endocytosis-related genes contribute to cancer cells’ resistance to carfilzomib. **A,** Cytoscape–generated ClueGO analysis of molecular pathways regulated by carfilzomib treatment (pathways association FDR<0.05, larger symbol size FDR<0.01), based on the proteomics data in Fig. 1. Colors, shapes and colored captions represent molecular pathways specific to cancer (red), normal fibroblast (dark blue), and multiple myeloma (magenta) cell lines, pathways common to two of them (dark yellow, orange and light blue), or all studied cell types (light green). Black captions show functional gene groups within the pathways (indicated with arrows) selected for siRNA vulnerability mini-screen. See Table S4 for details. **B, C,** Cancer cells and normal fibroblasts were transfected with a mixture of two siRNAs targeting genes within the indicated functional groups (all gene names listed in the Table S4) and treated with carfilzomib (CFZ) or vehicle (DMSO). siRNA targeting *PSMA2* gene served as a control targeting directly the proteasome bounce-back response. Viability was measured 24h after treatment using the ATP-lite luminescence assay (Methods). Panel **B** shows mean values for all tested cancer and normal fibroblast cell lines, panel **C** presents the viability of individual cell lines. Data in the heat map in **B** are means of n=6 for cancer and n=2 for normal fibroblasts, in **C** are means of n=2 for each cell line. Two-way ANOVA with Dunnett’s correction: *p < 0.05, **p < 0.01, ***p < 0.001 vs the CFZ-treated siRNA negative control (CTRL). **D,** Viability of the cancer cells transfected with an indicated combination or individual siRNAs targeting genes in the functional groups performing best in siRNA mini-screen (B,C) – molecular chaperones, autophagy or endocytosis - treated with CFZ or DMSO for 48h. Mix of siRNAs contains all 3 or 4 siRNAs used separately within a given functional group. Graphs present means of n=10, error bars represent SEM. Two-way ANOVA with Dunnett’s correction for the siRNA vs. the CFZ-treated siRNA negative control (CTRL) *p < 0.05, **p < 0.01, ***p < 0.001.

In the screen, the siRNAs targeting the chosen twelve functional protein groups were first used in pairs or trios (Table S4) to compare the viability of cancer cell lines to the normal fibroblasts (Fig. 2B, C). The siRNA sets targeting autophagy, molecular chaperones, and endocytosis were the most significantly decreasing viability of six cancer cell lines of lung, colon, and pancreatic cancer upon treatment with carfilzomib – either on average (Fig. 2B) or in the majority of the individual cell lines (Fig. 2C). The same siRNA sets were not decreasing cell viability significantly when supplementing the proteasome inhibition in the normal fibroblasts (Fig. 2B, C). This selected the targeted proteins in autophagy, molecular chaperones, and endocytosis pathways as the primary candidates for further testing as vulnerabilities in carfilzomib-treated cancer cells. We then tested how depletion of these proteins individually affects the viability of cancer cells *vs.* normal fibroblasts. HSP70 family proteins HSPA1A/B and HSP40 protein DNAJB1 in molecular chaperones, Cathepsins A and D in autophagy, and CHMP5 and RAB11A in endocytosis were the proteins whose depletion most significantly affected the survival of the six tested cancer cell lines combined with carfilzomib (Fig. 2D, Fig. S2B, C). We concluded that targeting these proteins and pathways could potentially increase the anti-cancer effect of proteasome inhibition in the therapeutic experimental setups.

### Inhibition of HSP70 proteins is efficient in the specific killing of cancer cells with the inhibited proteasome

We tested which of the currently available therapeutic inhibitors targeting autophagy, endocytosis, and molecular chaperone pathways are most efficient in killing the cancer cell lines with the inhibited proteasome, in comparison with the normal fibroblasts. Bafilomycin A1 (the inhibitor of autophagy), Hydroxychloroquine (the drug interfering with autophagy and endocytosis), or 17-AAG (the inhibitor of HSP90 family of molecular chaperones) were all significantly increasing the efficiency of carfilzomib in killing lung, colon, and pancreatic cell lines, however, they were similarly efficient in the killing of the normal cells (Fig. 3A). We then tested three different HSP70 family experimental inhibitors – VER-155008, JG98, and MAL3-101 ^39–41^. The results indicated that while MAL3-101 was toxic to normal fibroblasts (Fig. S3A), the two former inhibitors, especially VER-155008 at the lower of the used concentrations, were efficient in killing cancer lines and not the normal fibroblasts (Fig. 3B). To confirm this result in a more heterogeneous, patient-derived *in vitro* model, we used patient-matched tumor/normal organoid culture pairs, five in the colon and four in pancreatic cancer (Fig. S3B, C). Treatment of organoids with combinations of carfilzomib and VER-155008 or JG98 showed that the normal margin tissue from the same individuals is significantly less sensitive to the used drug combinations than the tumor-derived tissue and confirmed that the use of HSP70 inhibitors is augmenting the carfilzomib-dependent decrease of the viability in cancer-derived cells (Fig. 3C, D). A dispersion of organoids’ spheroid 3D structures was visible under the combinational treatment only in the cancer organoids compared to the normal tissue margin-derived organoids (Fig. 3E and Fig. S3D, E). These results prompted us to test the HSP70 inhibitors - VER-155008 or JG98 with carfilzomib *in vivo*, in subcutaneous xenografts of the cancer cell lines. First, we used the colon cancer DLD-1 cell line, where we tested which of the two HSP70 inhibitors was more efficient and less toxic to mice. In this experiment, JG98 was less efficient than VER-155008 in increasing carfilzomib’s effect on slowing down the xenograft growth (Fig. 3F, G), but it was also notably toxic to the mice – especially combined with carfilzomib causing diarrhea, mouse belly swelling, and the body mass decrease (Table S5). VER-155008 significantly increased the therapeutic effect of carfilzomib and was well tolerated by the animals (Fig. 3F, Table S6). The combination of VER-155008 and carfilzomib was also efficient in significantly slowing down the growth of the subcutaneous xenografts derived from the pancreatic cancer PANC-1 cells (Fig. 3F) and the lung cancer H-23 cells (Fig. 3G).

**Figure 3.**
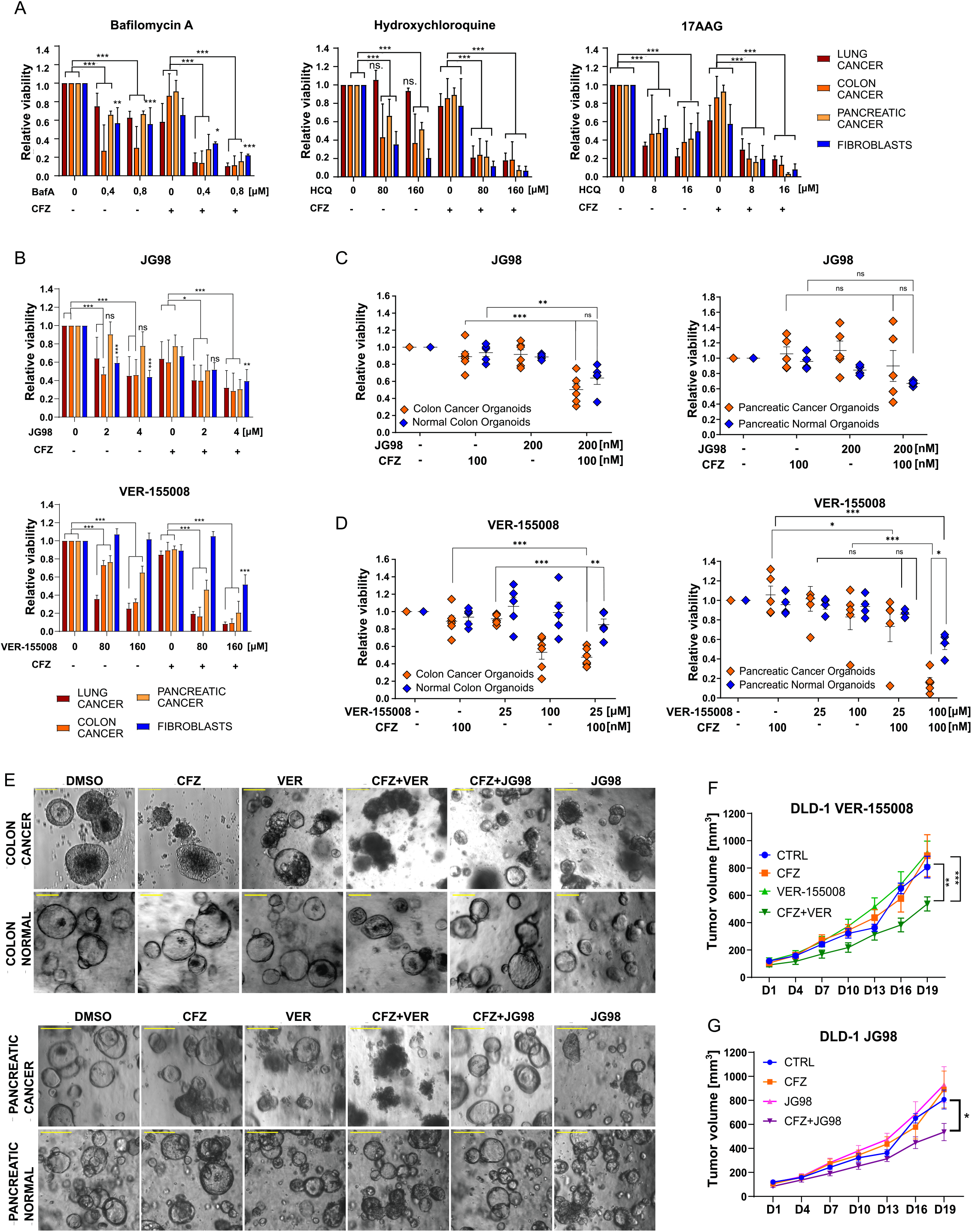
HSP70 inhibitors VER-155008 and JG98 specifically reduce cancer resistance to the proteasome inhibitor Carfilzomib. **A,** Viability of cancer cell lines (H-23, H1299 – lung; PANC-1, CAPAN-2 – pancreatic; RKO, DLD-1 – colon) and normal fibroblasts treated with carfilzomib (CFZ), HSP90 inhibitor 17AAG, and autophagy inhibitors bafilomycin A (BafA) and hydroxychloroquine (HCQ). Viability was measured with ATPlite after 48h of treatment with indicated drug concentrations. Error bars represent SEM. 2-way ANOVA with Tukey’s correction, *p < 0.05; **p < 0.01; ***p < 0.001; **B,** Viability of cancer cell lines (H-23, H1299 – lung; PANC-1, CAPAN-2 – pancreatic; RKO, DLD-1 – colon) and normal fibroblasts treated with carfilzomib (CFZ) and HSP70 inhibitors JG98 and VER-155008. Viability was measured with ATPlite after 48h of treatment with indicated drug concentrations. Bars represent mean of n=6. Error bars represent SEM. Two-way ANOVA with Tukey’s correction, *p < 0.05; **p < 0.01; ***p < 0.001. **C, D,** Viability of colon and pancreatic organoids derived from tumor and normal patient’s tissues. Organoids were treated with carfilzomib, VER-155008, and JG98 in indicated concentrations. Viability measured with ATPlite after 24h. Two-way ANOVA with Sidak’s correction, *p < 0.05; **p < 0.01; ***p < 0.001. **E,** Phase-contrast microscopy of colon and pancreatic organoids derived from tumor and normal patient’s tissues. Representative pictures of the organoid culture morphology, post treatment with proteasome and HSP70 inhibitors for viability test. Scale bar represents 200 µm. Corresponding fluorescent microscopy pictures of pancreatic, colon, and normal organoids are included in supplementary figure (Fig. S3 D, E) **F, G,** The size of xenografts formed in mice from DLD-1 cancer cells. Indicated drugs were administered intraperitoneally: DMSO – control group, carfilzomib (CFZ) 4 mg/kg, VER-155008 35 mg/kg, JG98 4 mg/kg, VER+CFZ 35 mg/kg + 4 mg/kg, JG98+CFZ 4 mg/kg + 4mg/kg, diluted in 200µl PBS-5% Tween-80 every other day. N = 6 animals in every group. Mixed-effect analysis with Sidak’s multiple comparisons test.

We concluded that the inhibition of HSP70 chaperones, especially with the inhibitor VER-155008, is the effective mean of specifically targeting cancer cells with the inhibited proteasome and not the normal cells. This prompted us to further investigate the mechanism of HSP70 proteins’ involvement in the response to the proteasome inhibition in cancer cells and to look for possibilities for further enhancement of proteasome and HSP70 targeting.

### High HSP70 level is associated with the resistance to the proteasome inhibition and low proteasome activity in neoplastic cells

To validate the response of proteins representing pathways important to the survival of the neoplastic cells on the inhibition of the proteasome machinery, we first compared their level changes upon treatment with carfilzomib or siRNA targeting PSMA2 20S proteasome core subunit essential for all the basic proteolytic activates of the 26S proteasome ^17^. Both carfilzomib and *PSMA2* siRNA resulted in the strongest and the most consistent upregulation of HSPA1A/B proteins among other molecular chaperones (Fig. 4A, B) chosen from the proteins significantly changing levels in the proteomics analysis and participating in the pathways enriched with the chaperone proteins (Fig. 1G, Fig. 2A, Table S4). Also, none of the tested proteins from the autophagy and endocytosis pathways was upregulated as consistently among cancer and normal cell lines as HSPA1A/B, on the proteasome inhibition (Fig. S4 A, B).

**Figure 4.**
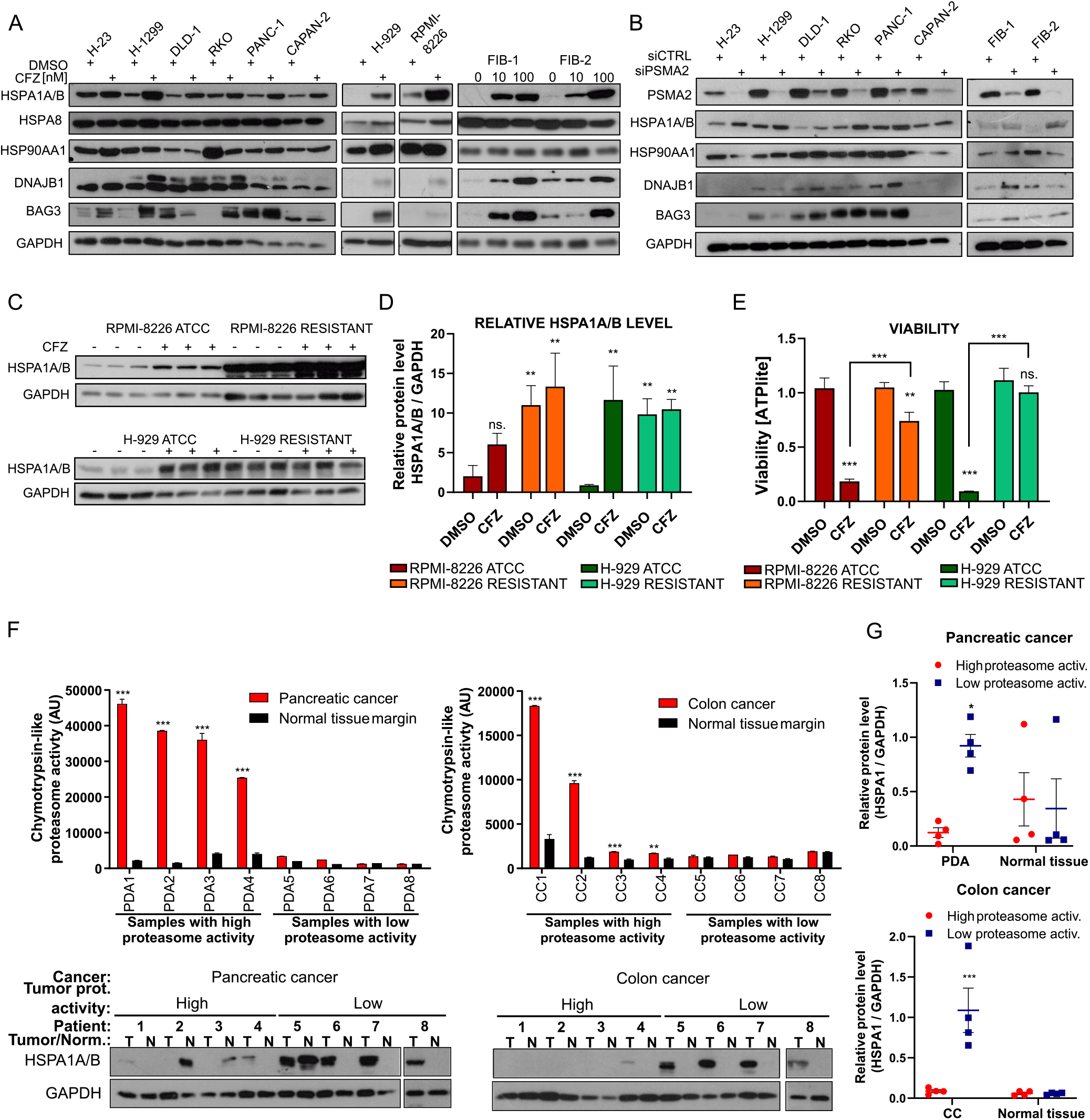
HSPA1A/B is upregulated under lowered proteasome activity conditions in human neoplasias. **A,** Molecular chaperone protein levels in cancer, multiple myeloma, and normal fibroblast cell lines treated with carfilzomib for 24h (Concentrations as in Figure 1E unless stated otherwise) determined by western blot **B,** Indicated molecular chaperone levels in cancer and normal fibroblasts transfected with siRNA targeting the proteasome subunit PSMA2, essential for the proteasome activity. **C,** H929 and RPMI-8226 multiple myeloma cell lines were selected cultured under increasing carfilzomib concentrations (0,1 nM – 30 nM) over 3 months’ time course to obtain cells resistant to the proteasome inhibitor. Western blot presents chaperons level and PARP cleavage in primary (ATCC) and resistant (RES) cell lines treated with carfilzomib for 24h. **D,** Level of HSPA1A/B protein measured by densitometry relatively to GAPDH in the western blot in C. **E,** Viability of primary and resistant multiple myeloma cell lines after treatment with carfilzomib, measured by the ATP-lite assay. **F,** Chymotrypsin-like proteasome activity in 8 pancreatic (PDA) and 8 colon cancer (CC) patient tissue samples, and corresponding normal tissue samples from the healthy margin of the same patients. The samples were designated “high proteasome activity” if the chymotrypsin-like proteasome activity in cancer samples was significantly higher than in the corresponding normal tissue sample; western blot shows HSPA1A/B and GAPDH protein levels in the samples. **G,** Relative HSPA1A/B protein levels normalized to GAPDH protein levels in “high” and “ low” proteasome activity sample groups from the western blot in F, in cancer and normal margin tissues; **E, F,** Bars are means of n=3 measurements with SD, test: two-way ANOVA with Bonferroni’s correction, *p < 0.05; **p < 0.01; ***p < 0.001; **G,** Horizontal, colored lines are means of n=4 plotted values with SEM, test: two-way ANOVA with Sidak’s correction, *p < 0.05; ***p < 0.001;

Multiple myeloma cell lines can be selected to resist the proteasome inhibitors, which is a clinically relevant problem ^10^. We tested if the resistance in H-929 and RPMI-8226 cell lines correlates with the increased basal HSPA1A/B protein levels. Resistant cell lines were obtained by a long-term adaptation to gradually increasing concentrations of carfilzomib starting at 0.1 nM. Indeed, the basal and carfilzomib-induced levels of HSPA1A/B in the obtained carfilzomib-resistant H-929 and RPMI-8226 cell lines, able to proliferate in up to 18 nM and 24 nM carfilzomib respectively, were higher than in the parental ATCC-derived cells (Fig. 4C, D). The viability of the carfilzomib-resistant cell lines strongly increased compared to the parental cell lines (Fig. 4E). This result suggested that in cancers, similarly to the carfilzomib-resistant multiple myeloma cells, a constantly high level of HSPA1A/B proteins may be present. To validate this hypothesis, we analyzed patient-derived tumor and normal tissue margin sample pairs in pancreatic and colon cancers (Fig. S4D). From deep-frozen tissue fragments available in our biobank we chose four sample pairs which upon a protein extraction showed the most significantly higher chymotrypsin-like proteasome activity than the patient-matched normal tissue fragments, and the same number of sample pairs with no significant difference of the proteasome activity in tumor *vs.* normal tissue (“high” and “low” proteasome activity samples, Fig. 4F). In both – pancreatic and colon tumor tissues - the HSPA1A/B proteins’ levels were on average several-fold higher in the samples with the low, than in the samples with the high proteasome activity, while no significant difference was observed in the patient-matched normal tissues (Fig. 4F, G).

The above results validated the proteomics results indicating a strong and consistent increase in HSPA1A/B protein upon the proteasome inhibition and inferred that the high level of HSPA1A/B proteins is an intrinsic attribute of neoplastic cells with low proteasome activity.

### HSPA1A/B is controlled by transcription on the proteasome inhibition and its mRNA level is a risk factor for cancer patients with a low proteasome subunit mRNA level

To understand how HSPA1A/B level is controlled upon the proteasome inhibition we studied the relation between the proteomes and transcriptomes of the neoplastic cells treated with carfilzomib. We performed a total RNA sequencing in cancer and multiple myeloma cell lines treated for 24h with DMSO (control) and the same carfilzomib concentrations as indicated in Fig. 1E, resulting in 50% ±10% decrease in chymotrypsin-like proteasome activity. Hierarchical clustering by Euclidean distance of quantified, protein-coding mRNAs (Table S7) showed a similar pattern as in the case of proteomes in Figure 1F, with cancer and multiple myeloma cells clustering separately (Fig. 5A). We performed differential mRNA level analysis between DMSO and carfilzomib-treated samples in all tested cell lines (Table S7). Correlation coefficients by Pearson or Spearman analysis between quantified proteins and protein-coding mRNAs of log fold changes in DMSO *vs.* carfilzomib sample groups (performed only for ID-matched protein-mRNA pairs), averaged in multiple myeloma and cancer cell lines, showed low mRNA-protein change correlation but suggested higher transcriptional control of protein levels in cancer than in multiple myeloma (Fig. S5B). We then matched only significantly up or downregulated proteins (p<0.05) to mRNAs in multiple myeloma and cancer cells (FDR<0.05). This analysis showed that most proteins significantly changing levels do not match the mRNA significantly changing levels in the same direction, with more proteins upregulated in multiple myeloma than in cancer (Fig. 5B). However, all the major upregulated chaperone proteins from HSP90, HSP70, BAG, or DNAJA/B families were present among the upregulated proteins direction-matching their significantly upregulated mRNAs in cancer cell lines pooled together (each listed chaperone gene in at least one cancer cell line). Indeed, the HSPA1A/B mRNA levels validated in the cancer cell lines treated with carfilzomib were on average over 30 times increased when compared to DMSO controls (Fig. 5C). Similar, albeit less strong responses, were visible in cases of other carfilzomib-responding molecular chaperones and co-chaperones – HSP90AA1, BAG3, and DNAJB1 (Fig. 5C), which together with earlier proteome–transcriptome changes matching result (Fig. 5B), suggested that proteasome inhibition leads to a general transcriptional induction of molecular chaperones. HSF proteins are the main factors controlling molecular chaperone genes’ transcription in stress conditions ^42^. We tested how knock-down of HSF-1 and HSF-2 transcription factors affects the mRNA and protein levels of molecular chaperones and we found that they were downregulated upon concomitant silencing of *HSF1* and carfilzomib treatment (Fig. 5C, D) and less significantly upon the silencing of *HSF2* (Fig. S5C). The most strongly HSF-1-dependent tested molecular chaperone on proteasome inhibition was HSPA1A/B (Fig. 5C, D). HSF-1 itself was not significantly upregulated in the proteomes of cancer cells treated with carfilzomib (Table S3), while we detected increased HSF-1 phosphorylation in selected cancer cell lines with the inhibited proteasome (Fig. S5D) - the known mechanism of HSF-1 activation in stress conditions ^43^. To test if broader targeting of molecular chaperone response, in addition to HSP70, further increases the effect of proteasome inhibition in cancer cells, we supplemented the carfilzomib and VER-155008 combination with HSF1 silencing. This however resulted in a stronger decrease in viability of normal fibroblasts, compared to cancer cells (Fig. 5E). Similar was the effect of silencing of BAG3 co-chaperone (Fig. 5E). Thus we concluded that targeting HSP70 is a more specific mean of specifically decreasing the viability of cancer cells with the inhibited proteasome, than broader targeting of the chaperone response.

**Figure 5.**
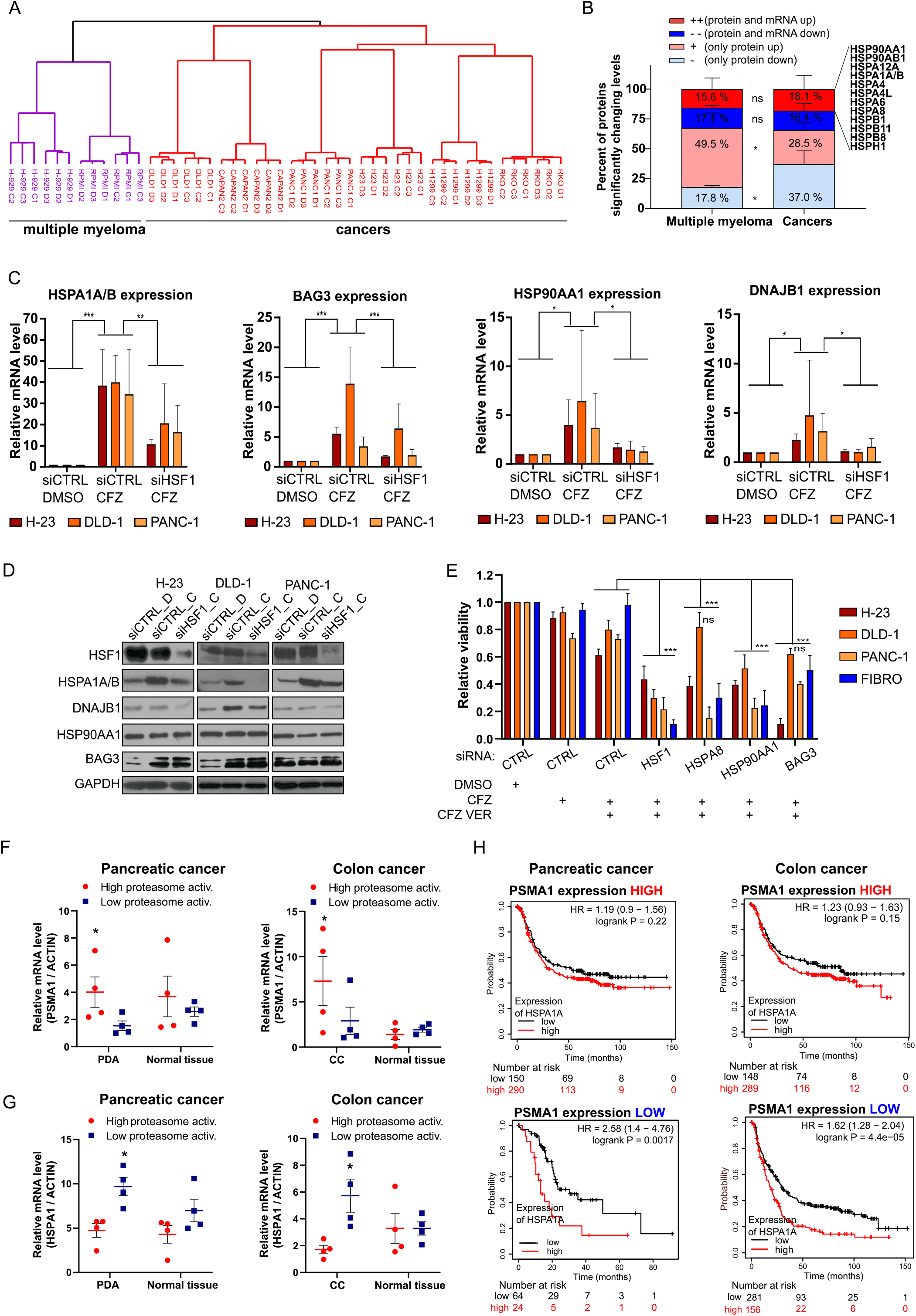
Transcriptome changes upon proteasome inhibition and HSF-1 dependent control of HSP70 expression in cancer cells. **A,** Hierarchical clustering (Euclidean distance) of transcriptomics results for 17311 mRNAs quantified in each of 36 transcriptomes from 8 indicated cell lines under control conditions (DMSO, D) or carfilzomib 24h treatment (C), each in n=3 biological replicates (full data in Table S5). Carfilzomib concentrations were as in Fig. 1E. **B,** Matching of the significantly up- or downregulated proteins (p<0.05, Table S3) to the mRNA significantly changing levels (FDR<0.05, Table S5) in cancer and multiple myeloma cell lines. Numbers of matched or unmatched proteins were transformed into percentages of all significant proteins in each cell line, means of these results were calculated (n=2 for multiple myeloma, n=6 for cancer cell lines; cell lines as in a), and plotted as stacked bars with SEMs. Two-way ANOVA with Holm-Sidak’s correction was used to compare corresponding protein groups between multiple myeloma and cancers, *p<0.05. A group of the key molecular chaperones is indicated, present in the upregulated protein fraction matching the significantly upregulated mRNA in cancers. **C,** Expression of HSPA1A/B, BAG3, HSP90AA1, DNAJB1 mRNA in the presence of carfilzomib (CFZ) and silenced HSF1 (siHSF1). H-23, DLD-1 and PANC-1 cancer cells were transfected with siHSF1 or siCTRL (negative control) and afterwards treated with carfilzomib (concentrations as in Fig. 1E). Expression of indicated genes was analyzed by qPCR. Bars represent mean of n=3 with SD. Two-way ANOVA with Tukey’s correction: *p<0.05; **p<0.01; ***p<0.001. **D,** Representative western blot analysis of HSF1 silencing and level of indicated chaperons in the presence of carfilzomib (CFZ) or DMSO control in the cell lines tested by qPCR (d). **E,** Viability of cancer and fibroblast cell lines treated with inhibitors and siRNAs. Cells transfected with indicated siRNAs were treated with CFZ (H-23 20nM, DLD-1 50nM, PANC-1 100nM, Fibroblasts 100nM), and VER-155008 40µM for 24h. Viability was measured with ATPlite. Two-way ANOVA with Dunnett’s correction: *p<0.05; **p<0.01; ***p<0.001; n=4 with SD. **F,** Proteasome subunit PSMA1 mRNA relative levels in “high” and “ low” proteasome activity sample groups from Fig. 4F, in cancer and normal margin tissues; **G,** HSPA1A/B mRNA relative levels in “high” and “ low” proteasome activity sample groups from Fig. 4F, in cancer and normal margin tissues; **H,** Association of the HSPA1A/B expression with patient’s survival in pancreatic and colon cancer patients datasets (Methods) in samples with high or low (above/below median) expression of PSMA1, representing the proteasome expression. Red curve – high level of HSPA1A/B; Black curve - low level of HSPA1A/B. HR – hazard ratio; log-rank P – log-rank test P value for the curves comparison. Numbers below graphs indicate number of patients at risk – total and at consecutive time points; n = 528 for pancreatic cancer, n = 874 for colon cancer

Since we found HSPA1A/B universally regulated by transcription in tested cancer cells with the inhibited proteasome (Fig. 5B-D), we investigated if HSPA1A/B mRNA level correlates with the proteasome activity and expression of the proteasome subunits. First, we used the patient-derived samples from Fig. 4F, with high or low proteasome activates in the pancreatic and colon tumor tissue. While higher mRNA levels of the representative proteasome core subunit genes – *PSMA1* or *PSMB5* – matched the high proteasome activity in the tumor samples (Fig. 5F, Fig. S5E), the *HSPA1A* gene had a reversed pattern of expression – significantly higher in the low proteasome activity tumor samples (Fig. 5G). This prompted us to test if in publically available datasets of cancer patients the proteasome expression level could affect how the *HSPA1A/B* expression correlates with patient survival. In the patients with the high *PSMA1* expression, which in our tests correlates with the high proteasome activity (Fig. 5F), the high level of *HSPA1A* expression had a less strong influence on the patient survival when compared with the patients with the low *PSMA1* expression. In the latter patients, the high *HSPA1A* expression correlated with a significantly worse prognosis than the low *HSPA1A* expression - in the colon and pancreatic cancers (Fig. 5H), as well as in other cancer types: lung, liver, head/neck, thyroid, ovarian, endometrial and in sarcomas (Fig. S5E).

We concluded that the main molecular chaperones, including HSPA1A/B, are primarily induced by the HSF-1-dependent transcriptional regulation across lung, colon, and pancreatic cell lines with the inhibited proteasome, while HSP70 remained more cancer-specific target upon proteasome inhibition than HSF1. This is in concordance with the association of the high *HSPA1A/B* mRNA levels with the low proteasome activity, which we observed in cancer patient tumor tissues, as well as with the worse prognosis in patients of multiple cancer datasets with the low proteasome subunit *PSMA1* expression levels.

### HSPA1A/B contributes to unfolded protein response and autophagy on the proteasome inhibition

The pathway analysis and the siRNA screen (Fig. 2A-C) indicated that autophagy, unfolded protein response (UPR) and endocytosis contribute to the resistance of cancer cells to the proteasome inhibition. We tested if these processes are a part of the HSP70 response network or are independent of HSP70 in the context of the carfilzomib treatment.

The UPR pathway is an important element of the cellular response to the accumulation of unfolded proteins in the ER, also in the presence of proteasome inhibitors ^44^. The main pro-survival UPR pathway is controlled by IRE1 receptor auto-phosphorylation and leads to an increased expression of chaperones and ERAD-related genes ^45^. To test if HSPA1A/B is involved in activation of this pathway on proteasome inhibition we analyzed the levels of total and phosphorylated (Ser 724) IRE1 in cancer cell lines transfected with siRNA targeting HSPA1A/B. As expected, treatment of the control cells with carfilzomib resulted in the increased phosphorylation of IRE1 (Fig. 6A). In cells with the silenced *HSPA1A/B,* the proteasome inhibitor did not induce the significant increase of phosphorylation of IRE1, measured as a proportion of phos-IRE1 to total IRE1 (Fig. 6B). To confirm the effect of HSPA1A/B on the IRE1 activation we tested its phosphorylation in the presence of VER-155008 alone and in the combination with carfilzomib. In all the tested cell lines the inhibitor of HSP70 significantly blocked the effect of carfilzomib on the increase of phosphorylated IRE1 (Fig. S6A, B), indicating the dependence of this process on HSP70.

**Figure 6.**
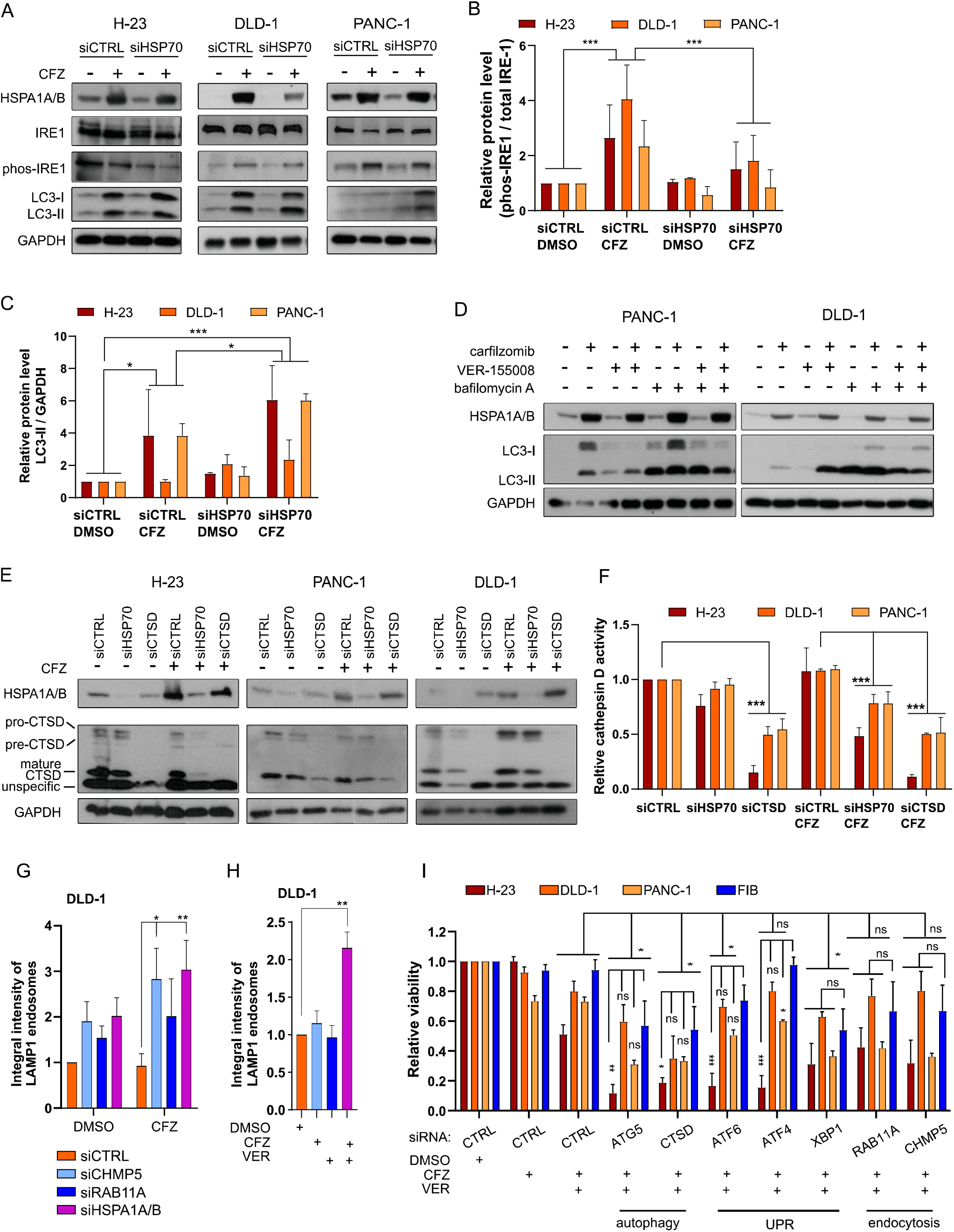
HSPA1A/B is involved in autophagy, unfolded protein response and endocytosis, contributing to cancer resistance to proteasome inhibitors. **A,** IRE-1 phosphorylation and LC3-I/LC3-II expression on HSPA1A/B (siHSP70) silencing and subsequent treatment with carfilzomib (CFZ) or control DMSO. Indicated cells were transfected twice day by day, treated with carfilzomib (concentrations as in Fig. 1F) a day after second transfection for 24 h. **B,** phospho-IRE-1 relative level on HSPA1A/B (siHSP70) silencing and subsequent treatment with carfilzomib. Bars represent mean of 2 or 3 independent experiments, based on western blot densitometry. Two-way ANOVA with Tukey’s correction, *p < 0.05; **p < 0.01; ***p < 0.001 **C,** LC3-II relative level on HSPA1A (siHSP70) silencing and subsequent treatment with carfilzomib. Bars represent mean of 2 or 3 independent experiments, based on western blot densitometry. Two-way ANOVA with Tukey’s correction, *p < 0.05; **p < 0.01; ***p < 0.001 **D,** LC3-I/LC3-II expression in indicated cell lines upon treatment with inhibitors: carfilzomib 100 nM, VER-155008 50 µM, bafilomycin A 20 nM for 24h. **E,** Cathepsin D (CTSD) level on HSPA1A/B (siHSP70) silencing and subsequent treatment with carfilzomib. siCTSD used as a control for cathepsin D activity test **(F)**. **F,** Cathepsin D activity on HSPA1A (siHSP70) silencing and subsequent treatment with carfilzomib. Two-way ANOVA with Dunnett’s correction, *p < 0.05; **p < 0.01; ***p < 0.001 **G,** Integral intensity of LAMP1 endosomes in DLD-1 colon cancer cell line transfected with indicated siRNAs and 24h post transfection treated with 50nM CFZ for another 24h. Two-way ANOVA with Dunnett’s correction, *p < 0.05; **p < 0.01, n=2 with SD, representative pictures Fig. S6E. **H,** Integral intensity of LAMP1 endosomes in DLD-1 colon cancer cell line treated for 24h with 50nM CFZ and 50µM VER-155008. One-way ANOVA with Bonferroni’s correction, *p < 0.05; **p < 0.01, n=2 with SD, representative pictures Fig. S6F. **I,** Viability of cancer and fibroblast cell lines treated with inhibitors and siRNAs. Cells transfected with indicated siRNAs were treated with CFZ (H-23 20nM, DLD-1 50nM, PANC-1 100nM, Fibroblasts 100nM), and VER-155008 40µM for 24h. Viability was measured with ATPlite. Two-way ANOVA with Dunnett’s correction: *p<0.05; **p<0.01; ***p<0.001; n=4 with SD.

It is known that proteasome inhibitors induce macroautophagy in multiple myeloma ^46^ and cancers ^21^. It was also demonstrated that HSPA1A/B is involved in the regulation of autophagy ^30,47^. To test the role of HSPA1A/B in the regulation of macroautophagy in cancer cell lines with the inhibited proteasome we compared the amount of LC3-II autophagosome marker ^48^ in relation to a housekeeping protein (GAPDH) in the cells transfected with a control or *HSPA1A/B*-targeting siRNA and treated subsequently with carfilzomib. We observed that in the cancer cells treated with carfilzomib levels of LC3-II were increased, while the concomitant silencing of *HSPA1A/B* and the inhibition of proteasome resulted in a further, significant accumulation of LC3-II (Fig. 6A, C). A similar effect on the level of LC3-II we found in the same cell lines treated with a combination of carfilzomib and VER-155008 (Fig. 6D). To discriminate if the LC3-II accumulation is a result of an autophagic flux increase or a block in the autophagosome accumulation, we treated the cells on inhibition of HSP70 and/or inhibition of the proteasome with bafilomycin A1, which blocks the function of lysosomes and their fusion with autophagosomes ^49^. In the case of carfilzomib and bafilomycin treatment the LC3-II accumulated further compared to carfilzomib alone, indicating an accumulation of autophagosomes and thus – the increase of the autophagic flux. In the case of the additional HSP70 inhibition by VER-155008, the accumulation of LC3-II was not further augmented by bafilomycin (Fig. 6D, Fig. S6C, D), indicating that HSP70 is maintaining the increase in an autophagic flux upon the proteasome inhibition.

Since the siRNA mini-screen showed the dependence of the cancer cells viability on lysosomal cathepsins during the proteasome inhibition (Fig. 2), we tested if in this condition HSPA1A/B affects the maturation and activity of the cathepsin D (CTSD). Silencing of *HSPA1A/B* upon the carfilzomib treatment caused a decrease in the mature form of CTSD detectable on western-blot (Fig. 6E) and a significant decrease of the CTSD activity (Fig. 6F). This result showed that HSPA1A/B affects the autophagic pathway induced by the proteasome inhibition on multiple stages – the autophagosomes accumulation and the cathepsin D maturation/activity.

Another process detected in the siRNA miniscreen (Fig. 2) to be important for the viability of cancer cells treated with carfilzomib was endocytosis, namely proteins involved in endosomal transport – RAB11A and an ESCRT-III complex CHMP5. While it is known that HSP70 is involved in early, clathrin-dependent endocytosis, the effect of the molecular chaperones as well as the proteasome inhibition on the subsequent endosomal transport has remained unknown. However, this phase of endocytic trafficking was shown to crosstalk with autophagy ^50^. We tested how proteasome inhibition and silencing of *RAB11A, CHMP5,* as well as *HSPA1A/B,* affect the accumulation of LAMP1-positive late endosomes, which mediate endocytic flux towards lysosomal degradation ^51^. In DLD1 cells, which responded most significantly of all the tested cancer cell lines, silencing of *RAB11A, CHMP5* and *HSPA1A/B* led to the accumulation of the LAMP1 endosomes (measured by fluorescence integral intensity of LAMP1 vesicles), which was further significantly increased upon treatment with the proteasome inhibitor (Fig. 6G, Fig S6E). This effect was mirrored by the dual treatment of cells with VER-155008 and carfilzomib (Fig. 6H, Fig S6F). Such an accumulation of LAMP1 endosomes could be indicative of improper endosome maturation and impeded endocytic flux ^52^, and the effect of HSP70 inhibition or silencing was comparable with depletion of endosomal traffic regulators – RAB11A and CHMP5.

To validate if HSP70 targeting complements or is redundant with simultaneous targeting of proteins involved in processes described in this section - autophagy, UPR, and endocytosis, we silenced their expression upon treatment of cancer and normal cells with carfilzomib and VER-155008 (Fig. 6I). This caused either a moderate additional decrease of viability in both cancer and normal cells or insignificant changes in the viability (Fig. 6I).

We concluded that HSPA1A/B functional effect on cancer cells upon the carfilzomib treatment extends to processes induced upon the proteasome inhibition specifically in cancer cells – UPR, endocytosis, and macroautophagy. This, however, results in redundancy of targeting these processes in addition to the proteasome and HSP70 in cancer cells.

### HSP70 chaperones the 26S proteasome and rescues its activity from the carfilzomib inhibition

Knowing that HSP70 family chaperones HSPA1A/B are strongly induced and important for cancer cells’ survival upon the proteasome inhibition (Figs. 1-5), we studied how mechanistically HSPA1A/B affects the processes involved in protein degradation and cell survival upon the carfilzomib treatment. First, we tested if HSPA1A/B depletion or overexpression affects directly the activity of the proteasome machinery in cancer and normal fibroblast cells. Chymotrypsin-like proteasome activity decreased insignificantly in control conditions on the silencing of *HSPA1A/B* with siRNA, while post-treatment with carfilzomib the silencing of *HSPA1A/B* caused an additional, significant decrease of the proteasome activity compared to the control siRNA, specifically in cancer cells (Fig. 7A). A similar effect was significant in the case of trypsin-like and caspase-like 26S proteasome activities, albeit as expected from previous reports ^12^ carfilzomib affected these proteolytic activities less strongly than the chymotrypsin-like activity (Fig. S7A, B). Additionally, treatment with VER-155008 significantly decreased the chymotrypsin-like proteasome activity upon the carfilzomib treatment in cancer cell lines and colon cancer organoids (Fig. S7C, D), while silencing of *HSP90AA1/B1* or *DNAJB1* genes had no significant effect (Fig. S7E). This suggested that the chaperone activity of HSPA1A/B could directly affect the proteasome in cancer cells. To validate this hypothesis we introduced to the study a dominant-negative K71S HSPA1A variant, unable to efficiently bind ATP and thus deprived of the chaperone activity ^53,54^. We overexpressed HA-tagged WT or K71S HSPA1A in control conditions and a silencing-rescue setup – upon *HSPA1A/B* silencing with an siRNA targeting the *HSPA1A* and *HSPA1B* mRNA 3’UTRs, in lung cell lines – H-23 and H-1299, suitable for efficient transient co-transfection experiments (Fig. 7B; Fig. S7F). The introduction of K71S HSPA1A in control conditions already affected negatively the chymotrypsin-like proteasome activity and the viability of the cells, while upon the carfilzomib treatment this effect was increased, and the overexpressed WT HSPA1A, but not the K71S variant, rescued the effect of silencing of the endogenous *HSPA1A/B* (Fig. 7C, D; Fig. S7G, H). This result supported the hypothesis that HSPA1A/B proteins may directly chaperone the 26S proteasome. Thus, we performed immunoprecipitation of the 26S proteasome core to detect interacting molecular chaperones. We detected HSPA1A/B but not HSP90AA1 interacting with the proteasome in three cancer cell lines, while in H-23 and PANC-1 cells the binding increased upon the carfilzomib treatment (Fig. 7E). To further validate the hypothesis, we reconstituted an *in vitro,* ATP-dependent, chaperoning system by using purified human HSPA1A, DNAJ1B, and HSP90AA1 proteins and the purified 26S proteasome as the potential chaperone client. The chaperones were first tested for their activity in an *in vitro* luciferase refolding assay ^55^, where the efficient refolding of the heat-inactivated luciferase requires the presence of all - HSP70 (in our case - HSPA1A), HSP40 (DNAJB1), and HSP90 (HSP90AA1) active proteins (Fig. S7I). When these chaperones were incubated with the 26S proteasome at 37°C it turned out that HSPA1A alone, and in the presence of DNAJB1, was significantly increasing the proteasome’s chymotrypsin-like activity, while HSP90AA1, DNAJB1 proteins alone, or the HSPA1A inactive variant K71S did not have this ability (Fig. 7F). Interestingly, the strongest increase in the proteasome activity was observed when the HSPA1A concentration was decreased and the HSP40 co-chaperone DNAJB1 was present – displaying a pattern of activity known from previous studies, where, in the presence of HSP40, increasing concentration of the HSP70 chaperone resulted in a loss of its optimal activity ^56^. The HSP70-dependent activation effect towards the purified 26S proteasome was inhibited by VER-155008, and the proteasome activity was inhibited by carfilzomib in all the tested conditions (Fig. 7F). However, the proteasome activity remained at significantly higher levels upon carfilzomib treatment in the presence of the HSPA1A-DNAJB1 chaperone setups optimal for supporting the 26S proteasome (Fig. 7F). This and the earlier results described in this section indicated that the HSP70 family chaperone proteins HSPA1A/B bind the 26S proteasome and decrease the inhibitory efficiency of carfilzomib *in vitro* and in cancer cells.

**Figure 7.**
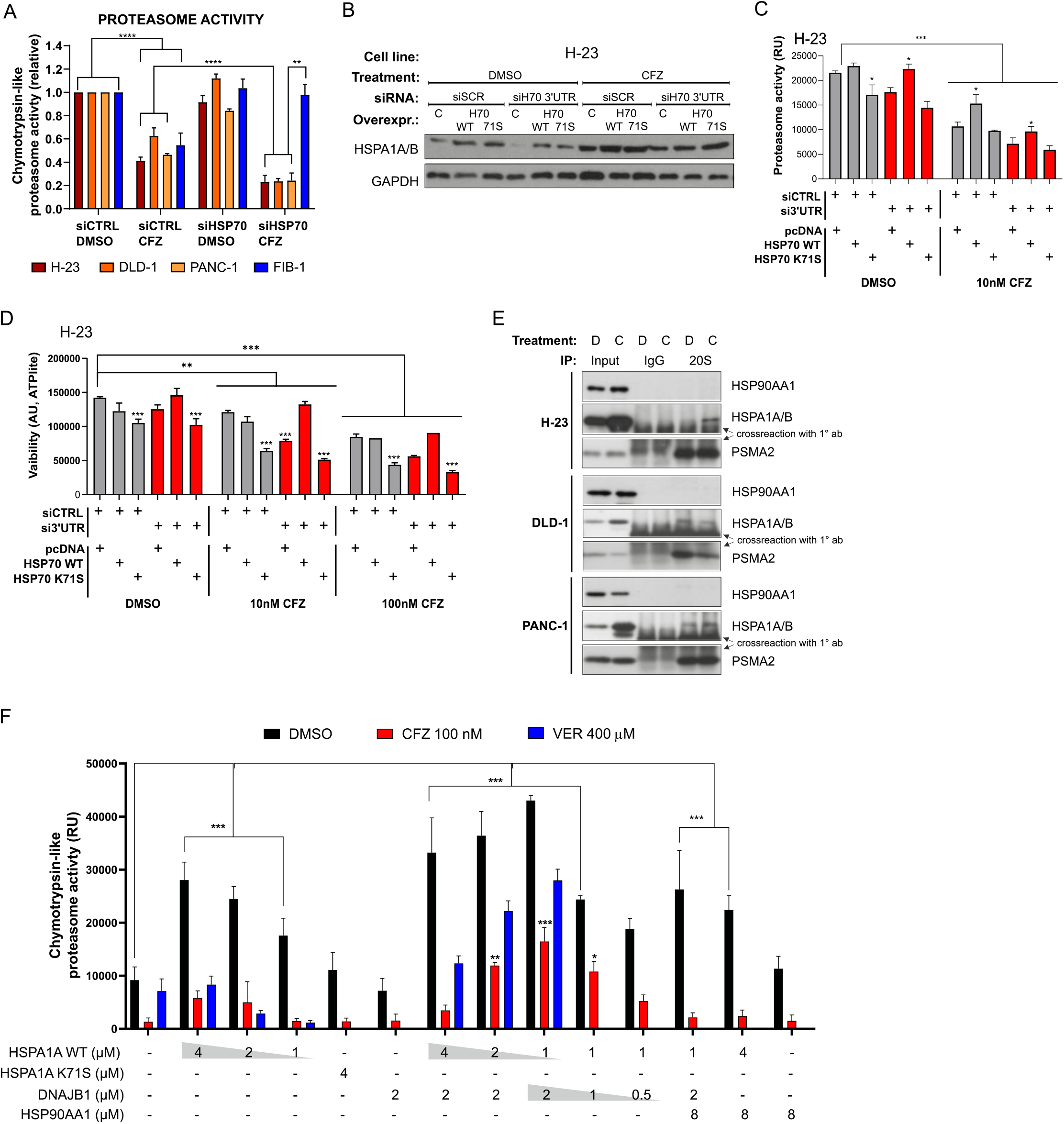
HSPA1A/B binds and chaperones the proteasome, increasing its activity and carfilzomib resistance in cancer cells. **A,** Chymotrypsin-like proteasome activity upon carfilzomib treatment (concentrations for as in Fig. 1E) and silencing of HSPA1A/B (siHSP70) in the indicated lung, colon and pancreatic cancer and normal fibroblast cell lines; siCTRL – negative siRNA control. Bars are means of n=5 with SD. **B,** Representative western blot demonstrating the silencing and the rescue overexpression of *HSPA1A* in H-23 lung cancer cell line. siCTRL – negative control; siH70 3’UTR – siRNA targeting 3’UTR of *HSPA1A/B* mRNA; H70 WT and H70 71S – vectors expressing HA-tagged wild type and K71S mutant of HSPA1A, respectively. Treatment with carfilzomib (CFZ) using concentrations as in Fig. 1E. **C,** Chymotrypsin-like proteasome activity in H-23 lung cancer cells upon silencing of *HSPA1A/B* and rescue overexpression of wild type (WT) or mutant (K71S) HSPA1A in the presence of indicated carfilzomib concentration. Bars are means of n=2 with SD. **D,** Viability of H-23 lung cancer cells upon silencing of HSPA1A/B and rescue overexpression of wild type (WT) or mutant (K71S) HSPA1A in the presence of indicated carfilzomib concentrations. Bars are means of n=3 with SD. **E,** Immunoprecipitation (IP) of the 20S proteasome with HSP70A1A/B (present) and HSP90AA1 (absent), along with indicated unspecific IgG controls and inputs from H-23, DLD-1 and PANC-1 cell lines, treated for 24h either with DMSO control (D) or carfilzomib (C) at 10 nM, 20 nM and 50 nM, respectively. Cross-reaction smears of the secondary detection antibody with the immunoprecipitation antibody are marked for clarity. **F,** Purified human 26S proteasome (at 5 nM, purified from HEK293 cells) chymotrypsin-like activity in the presence of indicated human molecular chaperones (recombinant, purified from *E.coli*), 100 nM carfilzomib and optionally – 400 µM VER-155008 HSP70 inhibitor. Bars are means of n=3, and for VER-155008 n=2, with SD. **A-G,** Two-way ANOVA with Tukey’s correction, *p < 0.05; **p < 0.01; ***p < 0.001;

### HSP70 does not affect the proteasome bounce-back effect which can be additionally targeted upon the proteasome inhibition

Since HSP70 increased the proteasome activity in cancer cells (Fig. 7), we additionally tested if HSPA1A/B affects the transcriptional bounce-back of the proteasome genes’ transcription ^16,17^, which could partially be responsible for the observed effect of HSPA1A/B on the proteasome. However, silencing of *HSPA1A/B* did not affect the compensatory bounce-back of two 26S proteasome genes representative for this process - *PSMA2* and *PSMC1* ^17^ - in cancer and normal cell lines (Fig. 8A and B). This remained in effect also for the treatment of cancer cells with carfilzomib and VER-155008 (Fig. S8A, B). Hence, we tested how the silencing of *NRF1* (*NFE2L1*) and *NRF2* (*NFE2L2*), the main transcription factors responsible for the bounce-back, affected the viability of cells treated with carfilzomib and VER-155008. The silencing of *NRF1* had a particularly significant effect in increasing the cytotoxicity of the drugs specifically in cancer cells, while silencing of both NRF1 and NRF2 was cytotoxic also to the normal fibroblasts (Fig. 8C).

**Figure 8.**
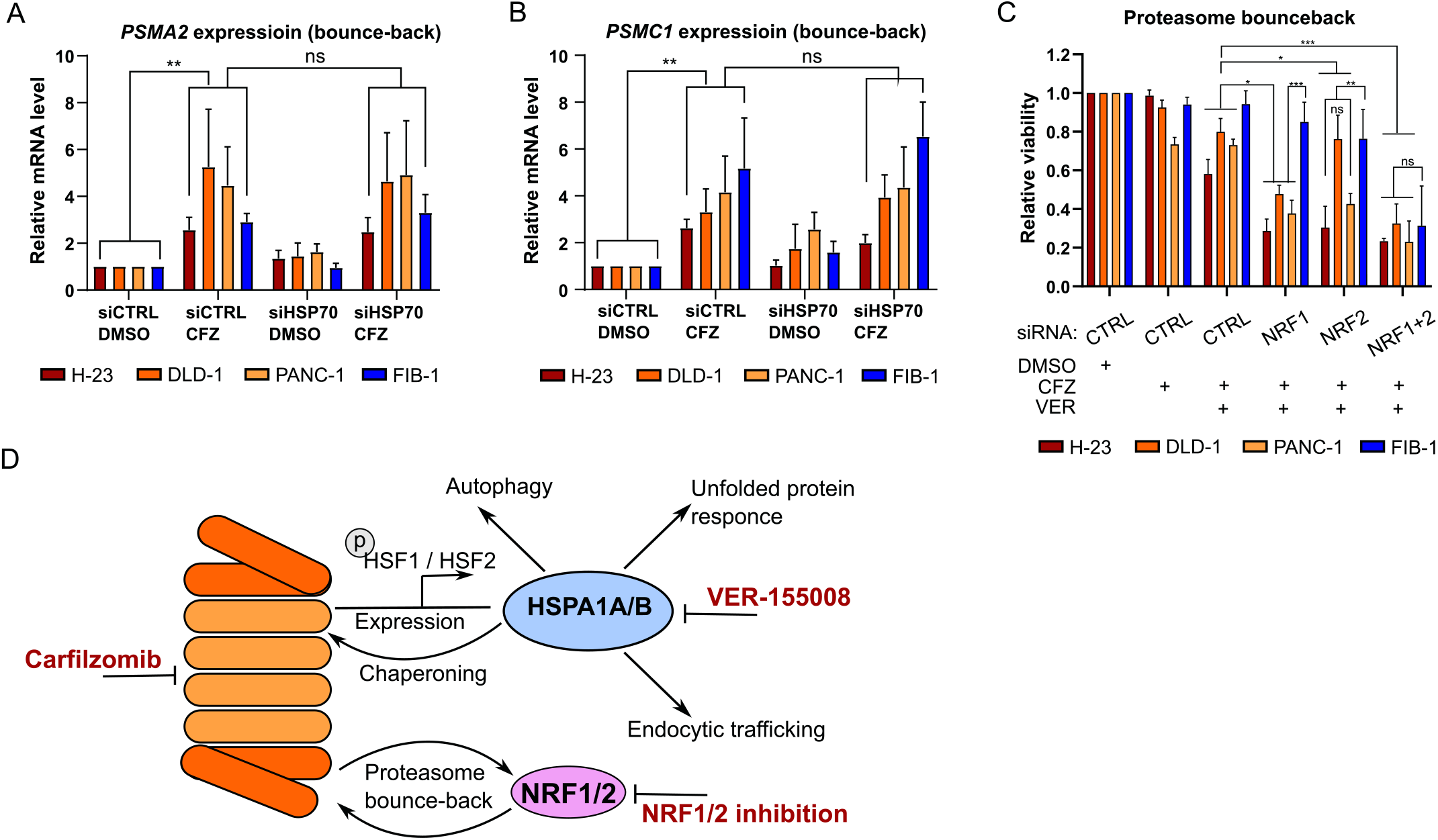
Proteasome bounce-back is independent on HSP70 and is a potential therapeutic target in combination with CFZ and VER-155008. **A, B,** Proteasome 20S core subunit *PSMA2* and 19S cap subunit *PSMC1* transcriptional bounce-back upon carfilzomib and silencing of HSPA1A/B (siHSP70) in the indicated lung (H-23), colon (DLD-1) and pancreatic (PANC-1) cancer and normal fibroblast (FIB-1) cell lines; siCTRL – negative control. Bars are means of n=4 with SD. C, Viability of cancer and fibroblast cell lines treated with inhibitors and siRNAs. Cells transfected with indicated siRNAs were treated with CFZ (H-23 20nM, DLD-1 50nM, PANC-1 100nM, Fibroblasts 100nM), and VER-155008 40µM for 24h. Viability was measured with ATPlite. Two-way ANOVA with Dunnett’s correction: *p<0.05; **p<0.01; ***p<0.001; n=4 with SD. **D,** Diagram summarizing the participation of HSP70 in the processes contributing to the resistance of cancer cells to proteasome inhibition.

Summarizing – targeting of HSP70 chaperones in cancer cells had the most significant, cancer-specific, cytotoxic effect, caused by simultaneous impact of HSP70 on multiple response mechanisms – such as autophagy, UPR, endocytosis, or directly - the proteasome activity. However, the NRF1/2-dependent proteasome expression bounce-back response bypassed the HSP70-dependence, rendering especially NRF1 an additional target in cancer cells with the inhibited proteasome and HSP70 (Fig. 8D).

## Discussion

In this study, we describe the proteasome inhibition compensatory responses by comparing protein landscape changes in normal fibroblasts, multiple myeloma, colon, lung, and pancreatic cancers cells treated with carfilzomib. This approach revealed the general proteasome inhibition response mechanisms, suggested by previous low-scale studies - proteasome subunit expression bounce-back ^16,26^, autophagy induction ^18,57^ or a molecular chaperone response ^27–30^ - as well as processes responding specifically in each cell type. In cancer cells, this included endocytosis, cell motility, mRNA splicing, or mitochondrial metabolism. Splicing and mitochondrial metabolism were previously found as mediators of the proteasome inhibitor resistance in proteomics studies in multiple myeloma cells ^58,59^. The possibility of comparing different neoplasia types in this study revealed that these processes are even more pronounced in cancer cells.

Our global proteomics experiments, unlike ubiquitination-focused studies ^60,61^, did not distinguish bona fide proteasome machinery substrates from compensators, and hence we used a vulnerability mini-screen to determine which of the found mechanisms contribute to the compensation of the carfilzomib treatment. The screen found as mediators of the compensatory response specific to cancer cells the molecular chaperones, autophagy, and endocytosis-related proteins. However, the drug tests in cancer cell lines, organoids, and *in vivo* xenografts as well as the low-scale validation pointed to the HSP70 family of proteins as the most cancer-specific component of the carfilzomib treatment resistance mechanism.

HSPA1A and HSPA1B are a duo of almost identical proteins, sharing a genomic locus ^36,62^, which concomitantly increase levels in response to various stressors, and are functionally interchangeable ^63^. Therefore we studied them as a single chaperone system in the inhibition/silencing experiments, while we used HSPA1A in the overexpression experiments and as a purified protein. HSPA1A/B along with other molecular chaperones and co-chaperones from HSP70, HSP90, BAG, and DNAJ families were reported to be induced on bortezomib treatment with involvement of HSF1 and HSF2 in multiple myeloma, breast, and cervical cancers or melanoma cell lines ^43,64,65^. We confirm this effect on carfilzomib treatment in a panel of lung, colon, and pancreatic cancer cells by transcriptome-proteome matching and low-scale validation. Consequently, we found that high levels of *HSPA1A* mRNA are associated with cancer *vs.* normal tissue and a bad prognosis in patients of multiple cancer types with a low proteasome subunits expression. However, silencing of *HSF1* along with inhibition of HSP70 and the proteasome turned out to significantly increase toxicity to normal cells.

The actual mechanistic purpose of the chaperone response in compensation of the proteasome inhibition has remained vaguely understood. The most often suggested hypothesis has been that the molecular chaperones facilitate folding and clearance of proteins accumulating upon proteasome inhibition, similar to the heat shock conditions ^66–69^. While our results are in concordance with such general effects, we show that HSPA1A/B specifically coordinates multiple independent routes of the proteasome inhibition response in cancer cells. HSP70 affects compensatory processes such as autophagy, unfolded protein response (UPR), and ESCRT-III-related endosomal transport, shown by us in the siRNA mini-screen and the functional experiments. These processes were earlier demonstrated in various experimental systems to respond to the downregulation of the proteasome activity ^18,19,23,70^, and separately selected aspects were shown to be dependent on HSP70 chaperones. ^47,71^. Our study links these findings by showing that in cancer cells HSPA1A/B inhibition significantly hinders the autophagic flux, maturation of cathepsin D, IRE1 phosphorylation, and progress of ESCRTIII-related endosomal transport exclusively upon the carfilzomib treatment – decreasing the efficiency of autophagy, UPR, and endocytosis responses to the proteasome inhibition. Thus, the dependence of these processes on HSP70 rendered their additional, direct downregulation redundant with the targeting of HSP70 and the proteasome in cancer cells.

We found that the binding and chaperoning of the 26S proteasome directly by HSP70 strongly decreases the efficiency of carfilzomib. Earlier, it was shown that individual chaperones may affect the assembly of the 26S proteasome, including HSP90 ^72^ or HSP70 upon the oxidative stress ^73^, with a HOP co-chaperone optionally involved in linking both systems ^74^. We demonstrate that the mature 26S proteasome is the HSP70 client, by reconstituting a functional *in vitro* chaperoning system. In this system, HSP70 can be stimulated for optimal efficiency by the HSP40 co-chaperone DNAJB1, while it does not require HSP90 proteins – reflected by a lack of HSP90 functional effect and binding to the proteasome in cancer cells. HSP90 often participates in the activation of intrinsically unstable substrates in their near-native states ^75^ or boosts the HSP70-mediated refolding of stress-unfolded substrates ^56^. This is a possible cause of the lack of effect of HSP90 on the assembled proteasomes which are relatively stable protein complexes ^76–78^. On the other hand, HSP70-DNAJ chaperone machinery possesses HSP90-independent activities, such as protein disaggregation ^79^, which could be in effect towards the proteasome in cancer cells. It can be speculated that the stress-induced nature of the HSPA1A/B action towards the proteasome makes it more indispensable in the cancer cells than in the normal fibroblasts (results in Fig. 7A) and allows cancer cells to maintain more proteasome activity under similar viability decrease as in the fibroblasts treated with carfilzomib (results in Fig. 1D). It has been described that an environment of highly stressed, rapidly evolving, and proliferating cancer cells is enforcing higher dependence of client proteins on HSP70 and HSP90 chaperone systems ^80,81^ – with the proteasome likely being another such example.

Importantly, we found that the HSP70 response network to proteasome inhibition does not include support of the proteasome subunit gene expression bounce-back, rendering unlikely the HSP70-dependent augmentation of the NRF1/NRF2 activity. As a consequence, additional targeting of NRF1 most significantly of all other tested targets enhanced the cytotoxic effect in cancer cells with the inhibited HSP70 and proteasome. Combinations of HSP70 inhibitors with bortezomib were suggested in multiple myeloma ^33^, melanoma ^31^, or bladder carcinoma ^82^, however, no clinical trials of any of these combinations have been carried out in cancer. Combining a direct or indirect NRF1 or NRF2 inhibition with inhibition of the proteasome was also elucidated in neoplasias ^83,84^, while not tested clinically in cancer. Especially methods of NRF1 targeting remain challenging and underdeveloped. Our results in lung, colon, and pancreatic cancers suggest that further preclinical and clinical investigations using a combination of carfilzomib, HSP70 inhibitors, and NRF1 inhibition methods are justified in an attempt to increase the efficacy of treatments in the most deadly cancer types.

## Methods

### Cell lines

NCI-H23 (H-23, RRID:CVCL_1547), NCI-H1299 (H-1299, RRID:CVCL_0060), DLD-1 (RRID:CVCL_0248) and CAPAN-2 (RRID:CVCL_0026) cell lines were cultured in RPMI medium (Gibco) supplemented with 10% FBS (Gibco) and Pen-Strep antibiotics (Gibco). RKO (RRID:CVCL_0504) and PANC-1 (RRID:CVCL_0480) cell lines were cultured in DMEM medium (Gibco) supplemented with 10% FBS and Pen-Strep antibiotics. Multiple myeloma cell lines RPMI-8226 (RRID:CVCL_0014) and NCI-H929 (H-929, RRID:CVCL_1600) were cultured in suspension, in RPMI medium supplemented with 10% FBS and Pen-Strep antibiotics (Gibco). Medium for H-929 cells was additionally supplemented with 0,05 mM of 2-mercaptoethanol (Bio-shop). All the above-mentioned cell lines were acquired commercially from ATCC repository and low passage numbers post acquisition (below 15) were used for the experiments in the manuscript.

Human primary fibroblasts FIB-1 and FIB-2 were cultured in DMEM medium (Gibco) supplemented with 10% FBS and Pen-Strep antibiotics. Fibroblasts were obtained and characterized at MMRC PAS, Warsaw, from skin biopsies of healthy control subjects, who provided informed consents based on bioethics committee approval 108/2017 and 203/2020 of the Central Clinical Hospital of Ministry of Interior and Administration in Warsaw ^85^.

All the cell lines were periodically controlled for mycoplasma infection (qPCR, all tests negative throughout the project) and none is present on the v11 of ICLAC register of misidentified cell lines.

### Mouse strains and animal care

NSG/J mice, purchased from The Jackson Laboratory (Bar Harbor, ME, USA), were maintained in an SPF facility under proper environmental conditions (20-24°C temperature, 40-60% humidity, and 12 h light cycle) with free access to water and food. The core of the breeding colony was a group of sister and brother (female homozygote × male homozygote) mated animals kept in Maria Sklodowska-Curie National Research Institute of Oncology bank of inbred strains.

### Human tissue samples, HE staining and tumor tissue identification

All experimental procedures using human tissue samples were conducted in accordance with an approval from ethical committee of Central Clinical Hospital of Ministry of Interior and Administration in Warsaw (109/2016 and 128/2018). All patients provided informed consent for the research use of their tissues. Colon cancer and pancreatic cancer tissues and adjacent normal colon and pancreatic tissues were obtained from patients undergoing surgical treatments at the Central Clinical Hospital of the Ministry of Interior and Administration in Warsaw and at the National Institute of Oncology in Warsaw. After preliminary histopathology assessment post-surgery the tissue fragments were either frozen and stored in a liquid nitrogen biobank for protein/RNA extraction (see appropriate sections) or kept in a culturing medium at 4°C for organoid cultures (see appropriate sections). Further histopathology microscopy assessment was used to confirm presence or absence of cancer tissue in the tissue fragments. Briefly, the tissue material from post-operative specimens was fixed in 10% buffered formalin, the samples were embedded in paraffin, sections of 4 µm thickness were stained with haematoxylin and eosin (HE) and observed/photographed using light microscopy. The histological type and the tumor grade were based on the WHO classification. The tumor stage was based upon the 8th edition of AJCC/UICC pTNM.

### Proteasome activity assay

Approximately 80% confluent cells were treated with concentrations of carfilzomib (Selleckchem) as in Fig. 1E unless indicated otherwise in the figures or a corresponding DMSO solvent volume as a control. Cells were washed with PBS, scraped from plates and lysed at 4°C in a lysis buffer containing 1% NP-40, 150mM NaCl, 50mM Tris-HCl at pH 8, and the cell remains were spun down. In the case of tissue samples, homogenization was performed using protein extraction beads (Diagenode) and sonication in the lysis buffer followed by centrifugation to remove solid remains (Diagenode Bioruptor Plus sonicator manufacturer’s protocol for protein extraction from tissue fragments). 50 µg of total protein extract was used per one measurement. Protein extracts in equal volumes were resuspended in 1x assay buffer containing 25mM HEPES pH 7.5, 0.5mM EDTA, 0.05% NP-40, 0.001% SDS to the volume of 90 µl per measurement and supplemented with 10 µl of 0.5mM proteasome substrates in the assay buffer: Substrate III (Suc-LLVY-AMC, chymotrypsin-like activity, Millipore), Substrate IV (Z-ARR-AMC, trypsin-like activity, Enzo) and Substrate II (Z-LLE-AMC, caspase-like activity, Enzo). Samples with substrates in 96-well black plates were incubated at 37°C for 2 h and measured using Tecan M1000 plate spectrofluorometer. Controls, sensitivity calibration and standard curves were made on the basis of recommendations of the 20S Proteasome Activity Assay Kit (Millipore).

### Cathepsin D activity assay

Cells were washed with PBS, scraped from plates and lysed at 4°C in a lysis buffer containing 1% NP-40, 150mM NaCl, 50mM Tris-HCl at pH 8, and the cell remains were spun down. 20 µg of total protein extract was used per one measurement. Protein extracts in equal volumes were resuspended in 1x assay buffer containing: HEPES 20mM pH 4, MgCl2 1mM, KOAc 137mM, NaCl 50mM, K2HPO4 50mM, EDTA 2mM, DTT 2mM to the volume of 90 µl per measurement and supplemented with 10 µl of 0.1mM cathepsin D fluorogenic substrate in assay buffer (Enzo). Samples with substrates in 96-well black plates were incubated at 37°C for 1h and measured using Tecan M1000 plate spectrofluorometer.

### Viability assays and drug tests

Cells were plated in 96-well white plates with clear bottom at 50% confluence for siRNA transfection (see ‘siRNA screen’ section for details) or 70% confluence for drugs test. VER-155008, hydroxychloroqiune, bafilomycin A (Tocris), MAL3-101 (A ChemTek, Inc), JG-98, 17-AAG, carfilzomib (Selleckchem) stock solutions were prepared in DMSO. The fresh medium with drugs at the concentrations indicated in the figures or DMSO solvent was added 24h and 48h after seeding. The viability was measured with ATPlite One Step reagent (Perkin Elmer) after 48h of the treatment. The results were reconfirmed with resazurin assay. The 10µl of resazurin solution in PBS (0.1mg/ml) was added directly to the cells growing in 200µl of culture medium and incubated for 3h in cell culture incubator. Afterward the fluorescence was measured (Ex/Em 530-560/590 nm) using Tecan M1000 plate spectrofluorometer.

### Proteomics analysis

Cells were lysed in 50mM Tris-HCl, pH 7.8 buffer containing 1% (w/v) SDS and 0.1M dithiothreitol, sonicated until the lysates were clear of long DNA and other visible cell debris (Diagenode Bioruptor Plus sonicator). Lysates were processed by the Multi-Enzyme Digestion Filter Aided Sample Preparation (MED FASP) protocol ^86^ with minor modifications ^17^. Briefly, proteins were digested overnight with endoproteinase LysC and then with trypsin for 3 h. The enzyme to protein ratio was 1:50. The total protein and peptide concentrations were determined by WF-assay ^87^. Aliquots containing 0.5 μg peptides were separated on a reverse phase C18 column and were analyzed on QExactive HF Mass Spectrometer (Thermo-Fisher Scientific) as described previously ^88^. Spectra were searched by MaxQuant software (https://maxquant.net/maxquant/) and the concentrations of proteins were assessed by the total protein approach using the raw protein intensities ^89^. Perseus software (https://maxquant.net/perseus/) was used to perform differential analysis, t-tests and assess p-value support of differences between protein concentrations in distinct experimental conditions.

### Pathway analysis

Proteins significantly changing levels (p<0.05) in the proteomics differential analysis in cancer, multiple myeloma and normal fibroblast cell lines were each fused in to signatures and filtered for duplicates. Such signatures were used in ClueGO ver. 2.5.8 ^90^ plug-in in Cytoscape ver. 3.8.2 (www.cytoscape.org) to associate proteins with molecular pathways. ClueGO settings were: All_Experimental evidence, GO Molecular Pathways/KEGG/WikiPathways ontologies, network specificity slider half way between Medium and Detailed settings, show only pathways with pV<0.05. Analyses performed for the separate - cancer, multiple myeloma and normal fibroblast signatures - were exported into tables and overlapped (Table S4) to determine pathways specific and common to the signatures. The graph in Fig. 2A was generated by performing the ClueGO pathway association with the same settings as above, using all three signatures as separate marker lists, and further node color/shape/link refinement in the Cytoscape environment according to signature overlap results in Table S4.

Ingenuity Pathway Analysis (IPA, Qiagen) was done by using protein lists significantly changing levels in each cell line (p<0.05 in the proteomics differential analysis) to perform core analyses. Results for canonical pathways were then overlapped in IPA separately for cancer, multiple myeloma and normal fibroblast cell lines and the overlaps were compared to find common and specific pathways to each cell line type.

### Plasmids

N- terminally HA-tagged WT and K71S HSP70 pCDNA 3.1 vectors were generated by cloning and subsequent site-directed mutagenesis at the Department of Molecular Biology at the International Institute of Molecular and Cell Biology in Warsaw ^54^. Expression vectors for chaperone protein overexpression in *E.coli* purification are described in the protein purification section.

### siRNA screen

siRNAs were purchased from QIAgen or Sigma Aldrich (company-validated, pre-designed sequences). Cells were plated in 96-well plates (white, transparent bottom) and transfected 2x at 0h and 24h with 20nM of siRNA (mixes or single) with GenMute transfection reagent (SignaGen) in cancer cell lines or Lipofectamine RNAiMAX (Invitrogen) in fibroblasts as in the manufacturer’s instructions. 48h after second transfection cells were treated with Carfilzomib or DMSO. Viability was measured after further 24h using ATPlite OneStep reagent (Perkin Elmer), according to the manufacturer’s instructions. The list of siRNAs is provided in the Table S8.

### Transfection

For siRNA transfections outside of the siRNA screen, all cells lines were transfected 2x at 0h and 24h with 20nM of indicated siRNAs using Lipofectamine RNAiMax (Invitrogen). After 24h post the second silencing, cells were processed. siRNA sequences are listed in Table S8.

H-1299 and H-23 cells were transfected with plasmids or co-transfected with plasmids and siRNAs using Lipofectamine 2000 (Invitrogen) as in the manufacturer’s instructions.

### Total RNA extraction and RT-qPCR

Total RNA was extracted from cell lines with QIAzol (Qiagen) following the manufacturer’s instructions, and form frozen tissue fragments - using QIAzol (Qiagen), RNA extraction beads (Diagenode) in a Diagenode Bioruptor Plus sonicator, according to a manufacturer’s RNA extraction protocol. RNA concentrations and quality was controlled using a NanoDrop spectrophotometer (Thermo). 1 µg microgram of total RNA was reverse-transcribed with NG dART RT kit (EURx). qPCR was performed using Sensitive RT HS-PCR Mix SYBR (A&A Biotechnology) on One Step Plus Real-Time PCR System (Applied Biosystems). The list of qPCR primers is provided in Table S8.

### RNA-sequencing

The cells were treated for 24h with Carfilzomib concentrations identical as for proteomics procedures in Fig.1E. Total RNA was extracted with QIAzol (Qiagen) following the manufacturer’s instructions, RNA quality was controlled using NanoDrop (Thermo) and RNA analyzer Experion (Biorad). Additional RNA quality check, library preparation (TruSeq Stranded TotalRNA, Illumina), sequencing (2x 100 bp, 100M reads; NovaSeq6000) and preliminary data quality check and analysis was performed by CeGaT GmbH. Quality assessment of the raw data including filtering and trimming was carried out using FastQC_v.0.11.9 (http://www.bioinformatics.babraham.ac.uk/projects/fastqc/) and the data was further processed using tool Trimmomatic_v.0.36 (http://www.usadellab.org/cms/?page=trimmomatic) for removal of low quality reads (phred-score ≤ 20 and length 30 bp, singletons discarded, reads with ambiguous bases ‘N’), trimming of bases from 5’/3’ end and adaptor sequences. Refined and filtered reads were further mapped to the reference human genome by Bowtie2 – used for indexing (http://bowtie-bio.sourceforge.net/bowtie2/), and HISAT2 – used for mapping (http://daehwankimlab.github.io/hisat2/). The successfully mapped reads were further processed to FeaturesCount R package to calculate the abundance of each transcript and extract quality score of mapped reads. Good quality scores were saved in a matrix form and were used as an input for determination of differentially expressed genes between control (DMSO) and carfilzomib treatment conditions by using DESeq2. The Relative Log Expression (RLE) method was used in DESeq2 to calculate normalization factors. Benjamini-Hochberg false discovery rate, FDR<0.05 was considered as statistically significant parameter of significantly differentially expressed genes. The count matrix genes were annotated with Ensembl BioMart to extract protein coding genes, which were used to generate principal component analysis (PCA) plot.

### Western blot analysis

Cell were lysed in a mild lysis buffer (150 mM NaCl, 1% NP-40, 50mM HEPES pH 8.0) supplemented with a HALT protease inhibitor cocktail (Thermo). Protein concentrations were determined using WF-assay ^87^. Lysates were incubated at 95°C in a Laemmli Sample Buffer, required protein amounts resolved by SDS-PAGE and transferred to nitrocellulose membrane (Millipore). Western blot analysis was performed according to standard procedures, using 5% fat-free milk in TBS-Tween20 0.1% to block and wash the membranes. The densitometry on WB bands was performed using ImageJ. The antibodies and concentrations used for WB are listed in Table S8.

### Human colon cancer and normal colon organoid cultures

Colon cancer and normal colon tissues were transported at 4°C in culturing medium w/o growth factors and processed within 18 h from resections. Protocol from ^91^ with small modifications was used to generate colon organoids. First, tissues were washed 10 times with ice-cold PBS, then minced using surgical scalpel. Minced tissues were incubated in digestion medium (Collagenase type II 5 mg/ml [GIBCO], Dispase 5 mg/ml [GIBCO], Y-27632 10.5 µM [Sigma], DNase I 10 µg/ml [Sigma], Advanced DMEM/F12 [GIBCO], HEPES 10 mM pH 7.5 [Invitrogen], GlutaMAX Supplement 1x [Invitrogen], Primocin 100 ug/ml [InvivoGen], Bovine Serum Albumin 0.1% [Sigma]) for 40 minutes at 37°C with rotation. Remaining undigested tissue fragments were allowed to settle to the bottom of the tube for 1 minute, then the supernatant was collected in a 15ml Falcon tube, centrifuged at 300 RCF for 5 minutes at 4°C. The cell pellet was embedded in a growth factor reduced Matrigel (Corning) or Cultrex Basement Membrane Extract type 2 (R&Dsystems). After 20 minutes incubation at 37°C, matrigel domes were covered with the culturing medium (Advanced DMEM/F12 [GIBCO], HEPES 10 mM pH 7.5 [Invitrogen], GlutaMAX Supplement 1x [Invitrogen], 10% R-spondin-1 conditioned medium ^92^, 50% Wnt-3A conditioned medium ^91^, N-acetylcysteine 1.25 mM [Sigma], Nicotinamide 10 mM [Sigma], B27 supplement 1x [GIBCO], Primocine 100 ug/ml [InvivoGen], murine Noggin 100 ng/ml [Peprotech], human EGF 50 ng/ml [Peprotech], human Gastrin I 10 nM [Tocris], Prostaglandin E2 10nM [Tocris], A83-01 500 nM [Tocris], Y-27632 10.5 uM [Sigma], SB202190 3 uM [Sigma]).

### Human pancreatic cancer and normal pancreatic organoid cultures

Pancreatic organoids were generated in a similar way to colon organoids based on ^93^, with addition of human FGF-10 100 ng/ml (Peprotech) and no SB202190 in the culturing medium. Prostaglandin E2 10nM (Tocris) was added only to medium for normal tissue organoids.

### Drug sensitivity assays in cell lines and organoids

Cell lines were treated by adding the drugs (or their solvents in equal volumes) at concentrations indicated in the figures to the standard growth media. Viability assay was performed after indicated time periods.

For organoids 96-well plates with clear bottom and white walls were used for testing. 33 µl of Matrigel or Basement Membrane Extract type 2 were added to each well. Plates were then centrifuged at 1000 RCF for 1 minute and placed in a 37°C, 5% CO_2_ incubator for 30 minutes. To each well a 100ul suspension of approximately 500 organoids in the culturing medium were added. Organoids were left for 24h in a 37°C, 5% CO_2_ incubator. Afterwards the drugs were added in the culturing medium. Organoids plates were incubated with drugs for 24h, then cell viability was measured using ATPlite Luminescence Assay (PerkinElmer).

### Immunofluorescence staining of LAMP1 endosomes

LAMP1 endosomes were detected by immunofluorescence staining using mouse anti-LAMP1 (H4A3) antibodies generated by J. Thomas August and E.K. Hildreth from the Developmental Studies Hybridoma Bank developed under the auspices of the NICHD and maintained by the University of Iowa, Department of Biology, Iowa City, Iowa, USA. DLD-1 cells were seeded on µClear 96-well plates (Greiner Bio-One, #655096) and transfected twice at 0h and 24h with 20 nM of indicated siRNAs using Lipofectamine RNAiMax (Invitrogen). After 24h following the second transfection, cells were transferred to ice and washed twice with ice-cold PBS and ice-cold 3% paraformaldehyde was added to cells for 15 min at RT. Then, cells were washed three times with PBS and stained directly or stored at 4°C. For staining, cells were incubated for 10 min with 0.1% (w/v) saponin (Sigma-Aldrich), 0.2% (w/v) fish gelatin (Sigma-Aldrich), and 5 mg/ml BSA (BioShop) in PBS. Afterward, cells were incubated with primary anti-LAMP1 antibodies (1:800) diluted in 0.01% (w/v) saponin and 0.2% fish gelatin in PBS for 1h. Then, cells were washed twice for 5min with 0.01% (w/v) saponin and 0.2% fish gelatin in PBS. Next, cells were incubated for 30min with donkey anti-mouse Alexa 488 secondary antibodies (Life Technologies, A21202) and DAPI (1µg/ml; Sigma-Aldrich). The images were obtained using Opera Phenix confocal microscope (PerkinElmer) with 40x/1.1 water immersion objective. Harmony software(version4.9; PerkinElmer) wasused for image acquisition and analysis. At least twenty 16-bit images with resolution 1080 x 1080 pixels were acquired per experimental condition. Flatfield correction (advanced algorithm from Harmony software) was applied to all images before analysis. The image analysis was performed using the spot detection function to detect mask for vesicles positive for LAMP1. Subsequently, the integral fluorescence of LAMP1 marker in the mask region was counted. Cell number was determined by nuclei detection using DAPI signal. All data were normalized to cell number. Pictures were assembled in Photoshop (Adobe) with only linear adjustments of contrast and brightness.

### Immunofluorescence staining of human colon and pancreatic organoids

Organoids cultured in a chamber slide (Ibidi) and incubated for 24h with the indicated drugs, were fixed with 4% PFA for 20 minutes, washed with PBS/Glycine solution, permeabilized with 0.5% TritonX-100/PBS for 10 minutes and blocked with 3% bovine serum albumin/PBS. Incubation with a primary antibody was done overnight at 4°C. Primary antibodies used: E-Cadherin (Cell Signaling Technology, 24E10) and Laminin-5 (Santa Cruz Biotechnology, P3H9-2). Incubation with a secondary antibody was done overnight at 4°C. Secondary antibodies used: Alexa Fluor 488 (Life Technologies, A11001) and Alexa Fluor 555 (Life Technologies, A21428). Incubation with 0.5 ng/ml DAPI in PBS for 10 minutes was used for staining of nuclei. Image acquisition was performed using Carl Zeiss LSM 780 confocal microscope.

### Kaplan-Meier estimator analysis

The analysis was performed in the KM-plotter (kmplot.com) using the mRNA gene-chip data for lung cancer and the pan-cancer RNA-seq data for other cancer types shown in the figures. The multiple genes option was used to filter the results by a median of *PSMA1* expression (above/below median – high/low expression) and patients survival was further stratified according to *HSPA1A* high/low expression. The log-rank test and hazard ratio (HR) estimation were performed by the KM-plotter tool to assess the effect of gene expression on the patient survival.

### Co-immunoprecipitation

Cell lines were treated with carfilzomib or DMSO as for proteomics analysis. After 24h formaldehyde cross-linking was performed according to the protocol from ^94^. Briefly, cells were treated with 0.4% formaldehyde solution in PBS for 7 minutes at room temperature (1ml of formaldehyde solution per 1×10^7 cells). Cells were then pelleted at 1800 RCF for 3 minutes at room temperature. Cell pellet was resuspended in 0.5ml ice-cold 1.25M glycine/PBS and centrifuged at 1800 RCF for 3 minutes at 4°C. Cells were lysed (150 mM NaCl, 50 mM Tris-HCl pH 7.4, NP40 1% and protease inhibitors [Thermo]) and immunoprecipitation was performed overnight at 4°C using Proteasome 20S core subunits polyclonal antibody (ENZO Life Sciences BML-PW8155) and IP buffer (137 mM NaCl, 50 mM Tris-HCl pH 7.4, NP40 1%, 2 mM EDTA pH 8 and 10% glycerol). Immunopellets were washed 3 times with the IP buffer without glycerol, and boiled in the 1x Laemmli sample buffer for 15 minutes. SDS-PAGE and western blot was performed. Detection of Hsp70, Hsp90 and 20S Proteasome was done using primary antibodies: HSP70 (ENZO Life Sciences, ADI-SPA-812-F), HSP90 (Cell Signalling, C45G5) and PSMA2 (Cell Signaling, 2455S). TidyBlot:HRP conjugated Western Blot Detection Reagent (Bio-Rad) was used in accordance with manufacturer’s protocol as the secondary antibody.

### Protein purification, luciferase refolding assay and *in vitro* 26S proteasome chaperoning reaction

Human HSPA1A (Hsp70) WT and K71S, HSP90AA1 (Hsp90α), DNAJB1 (Hdj1) were purified and tested by a luciferase refolding assay as described previously ^54,55^. Human 26S proteasome purified from HEK293 cells was purchased from Promega. The proteasome chaperoning reaction tests were carried out as follows: 5 nM of the purified proteasome and the molecular chaperone proteins (or a mass equivalent of BSA as the control) in concentrations indicated in the figure were added on ice to a final reaction volume of 10 µl with the reaction buffer containing: 10 mM Tris-HCl pH 7.5, 50 mM KCl, 3 mM MgCl_2_, 2 mM DTT, 5 mM ATP, ATP-regeneration system (0.3 units of creatine kinase, 150 mM of phosphocreatine; Roche) and optionally: DMSO (drug solvent control), carfilzomib or VER-155008. The reaction mixes were incubated for 30’ at 37°C. The proteasome activity assay followed by addition to each mix of 80 µl of the 1x assay buffer + 10 µl of 0.5mM proteasome chymotrypsin-like proteasome activity substrate and incubation for 1h at 37°C in a 96-well black plate before fluorescence measurements (see the Proteasome activity assay section).

### In vivo xenograft experiments

Experiments were carried out in the animal facilities of Maria Sklodowska-Curie National Research Institute of Oncology, in accordance with the protocols approved by the Second Local Ethics Committee for Animal Experimentation in Warsaw (decision no. WAW2/117/2020). Xenograft implantations were performed on 6–18 week-old mice in a separate operating room using aseptic procedures. To induce subcutaneous xenografts from cultured cell lines, 2 × 10^6^ of cancer cells (DLD-1, H-23 and PANC-1) suspended in 100µl PBS were injected subcutaneously into a flank of eight animals per group. Tumor diameters were measured weekly with a caliper until reaching a volume of 100 mm^3^, calculated using the following formula: (length × width × width)/2.

### Animal groups and drug administration

When all the tumors within a set exceeded the required volume the animals were assigned randomly to treated and control groups. Experiment 1, DLD-1: 7 x 6 animals; DMSO – control group, carfilzomib - CFZ (Selleckchem, 4 mg/kg), VER-155008 (Tocris, 35 mg/kg), JG98 (Selleckchem, 4 mg/kg), VER+CFZ (35 mg/kg + 4 mg/kg), JG98+CFZ (4 mg/kg + 4 mg/kg), were diluted in 200µl PBS-5%Tween-80 and injected intraperitoneally every other day. Experiment 2 and 3, PANC-1 2×7 animals and H-23 2×5 animals: DMSO – control group, VER+CFZ (35 mg/kg + 4 mg/kg). Mice were carefully observed for the appearance of signs of distress. Tumor diameters were measured every third or fourth day with a caliper. On day 21 or when a tumor volume exceed 1500 mm3, mice within a set were sacrificed. Blood was collected, and tumors were excised for histopathological and molecular examinations.

### Materials availability

This study did not generate new unique reagents.

### Statistical analysis

All data are represented as mean ± standard deviation (SD) or standard error of the mean (SEM). Figure legends contain information on independent biological replicates and statistical tests used. The statistical analysis was performed using GraphPad Prism 8.0.2. Proteomics, RNAseq and Pathway analysis sections contain details on statistical analysis used in large-scale and pathway analysis procedures.

## Data availability

Mass spectrometry: ProteomeXchange Consortium via the PRIDE partner repository https://www.ebi.ac.uk/pride/ with the dataset identifiers: PXD025364 (cancer cell lines), PXD027840 (normal fibroblast cell lines)

RNA sequencing: Gene Expression Omnibus (GEO), accession code GSE184029. https://www.ncbi.nlm.nih.gov/geo/query/acc.cgi?acc=GSE184029

## Acknowledgements

The authors thank prof. Maciej Żylicz for providing vectors and purified proteins obtained at the Department of Molecular Biology of the International Institute of Molecular and Cell Biology in Warsaw and prof. Cezary Żekanowski for sharing laboratory equipment at the Laboratory of Neurogenetics, Mossakowski Medical Research Institute PAS. We thank all cancer patients from hospitals in Warsaw, Poland, who provided consent to use their tissue samples in the manuscript as a part of the “Multi-onko-mapa” study (109/2016). This research was funded by the National Science Center, Poland grants Opus 2017/25/B/NZ5/01343 (D.W.), Miniatura 2020/04/X/NZ5/01259 (M.O.) and EU H2020 Marie Curie Individual Fellowship 795441 (D.W.).

## Author contributions

Conceptualization, D.W., M.O. and M.G.; Methodology, D.W. M.O., M.G, A.J. and J.R.W.; Investigation, D.W. M.O., M.G, A.J., K.J., J.L., M.C., M.Miaczynska., K.Z., J.R.W.; Data Curation, D.W. and A.J., Writing – Original Draft, D.W and M.O.; Writing – Review & Editing, D.W. and M.O.; Funding Acquisition, D.W. and M.O.; Resources, M.N-N., M.K., W.K., T.O., A.Z., M.Mikuła., and M.L.; Supervision, D.W.

## Declaration of interests

The authors declare no competing interests.

## Notes

### Competing Interest Statement

The authors have declared no competing interest.

