## Supplementary figures for "The response network of HSP70 defines vulnerabilities in cancer cells with the inhibited proteasome"

Figure S1

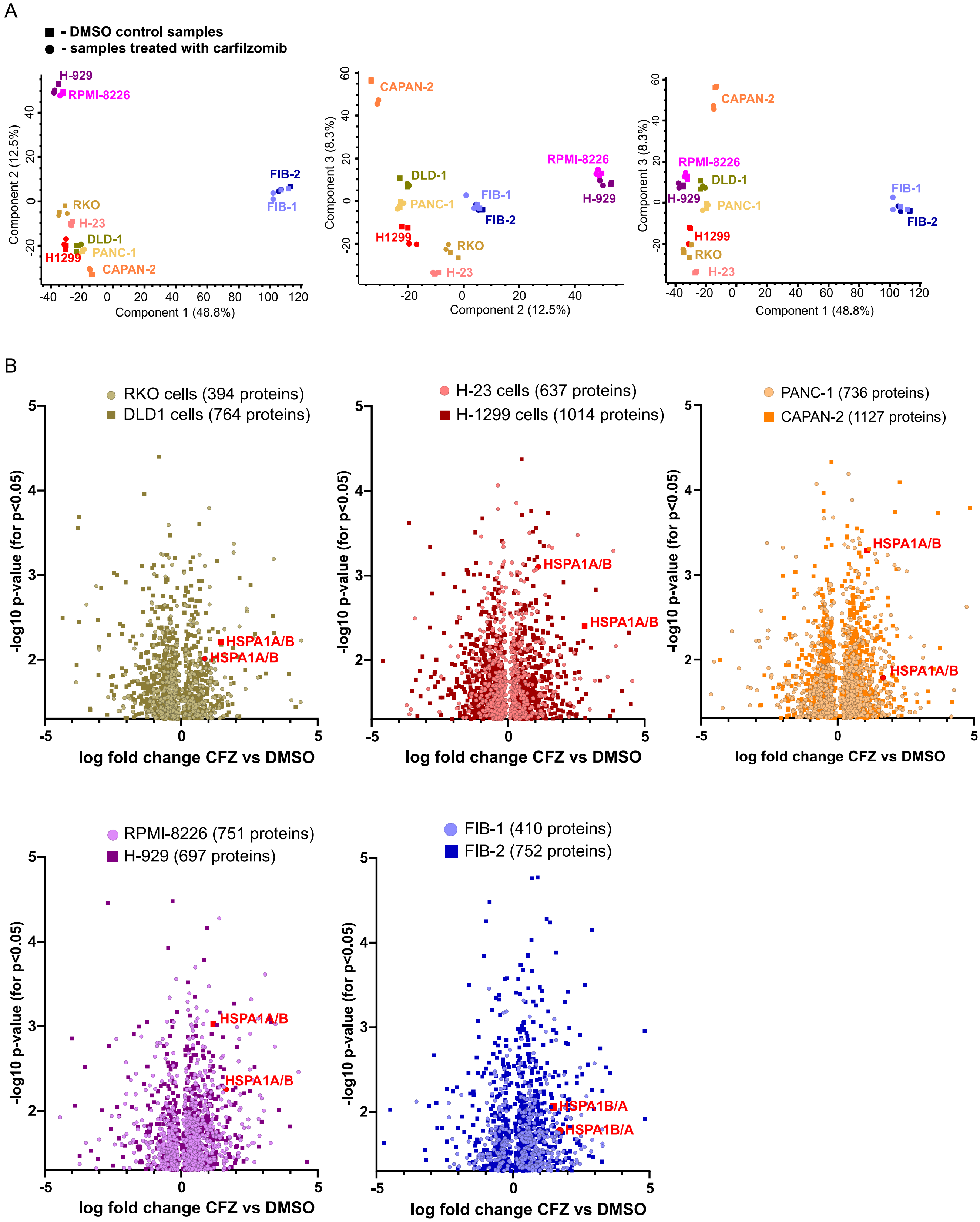
A
■ - DMSO control samples
● - samples treated with carfilzomib
H-929
RPMI-8226
FIB-2
FIB-1
RKO
H-23
DLD-1
PANC-1
H1299
CAPAN-2
Component 2 (12.5%)
Component 1 (48.8%)
Component 3 (8.3%)
Component 2 (12.5%)
Component 3 (8.3%)
Component 1 (48.8%)
B
RKO cells (394 proteins)
DLD1 cells (764 proteins)
H-23 cells (637 proteins)
H-1299 cells (1014 proteins)
PANC-1 (736 proteins)
CAPAN-2 (1127 proteins)
RPMI-8226 (751 proteins)
H-929 (697 proteins)
FIB-1 (410 proteins)
FIB-2 (752 proteins)
HSPA1A/B
HSPA1B/A
-log10 p-value (for p<0.05)
log fold change CFZ vs DMSO

**Figure S1**

**A,** PCA (principal component analysis) in proteomics samples for cell lines indicated in the graphs, under control conditions (DMSO) or carfilzomib treatment.

**B,** Volcano plots representing differential expression analysis results in the indicated cell lines (log fold change of carfilzomib (CFZ)-treated samples vs. DMSO controls was plotted against –log10 of p-values, for significant results of $p<0.05$). The plots are in pairs of cell lines – of colon, lung, pancreatic cancers, multiple myeloma and normal fibroblasts. HSPA1A/B protein, the only one significantly changing level (upregulated) in all studied cell lines, is marked in red.

Figure S2

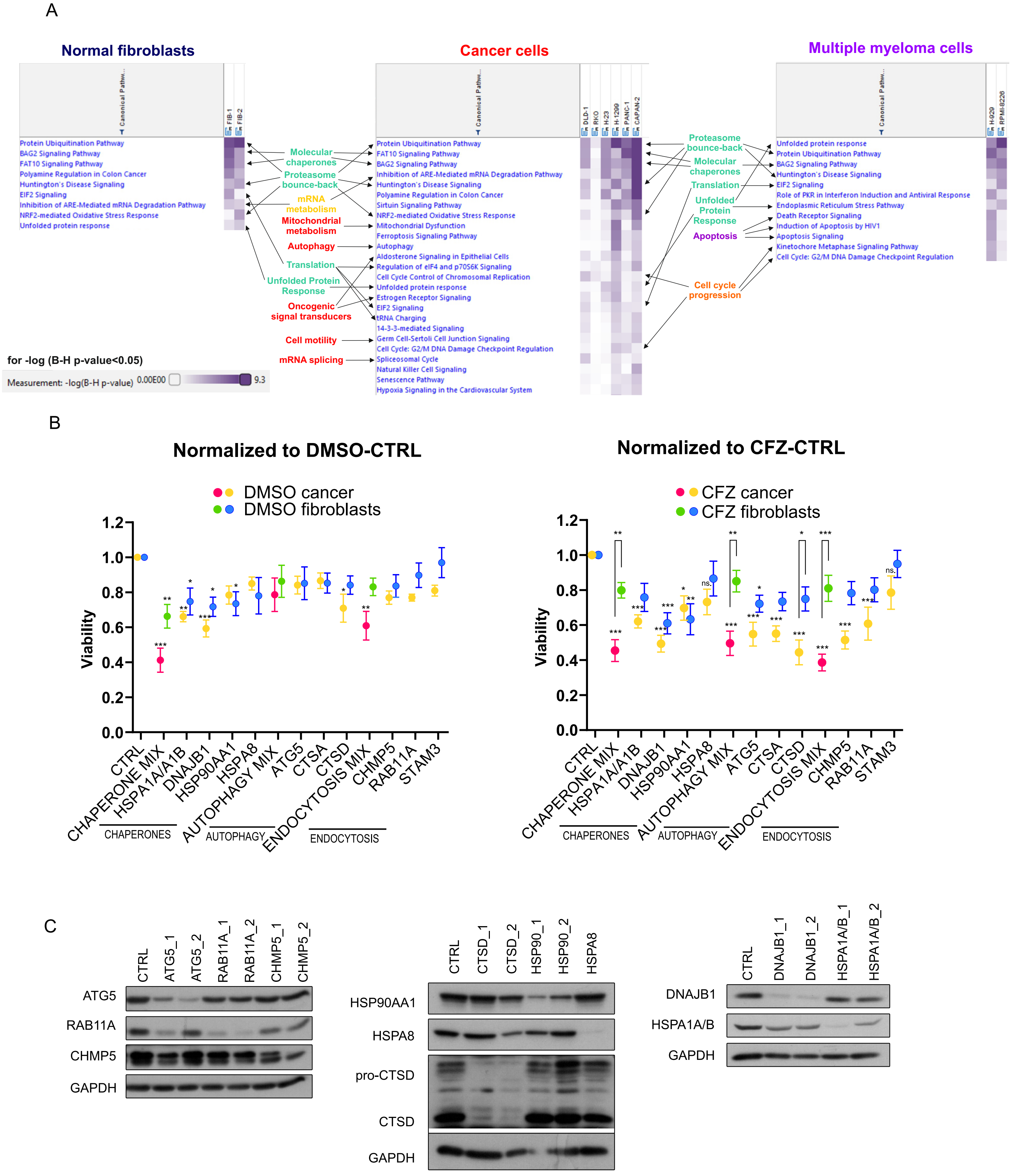

A
Normal fibroblasts
Cancer cells
Multiple myeloma cells
Canonical Pathw...
FIB-1
FIB-2
DLD-1
RKO
H-23
H-1299
PANC-1
CAPAN-2
H-929
RPMI-8226
Protein Ubiquitination Pathway
BAG2 Signaling Pathway
FAT10 Signaling Pathway
Polyamine Regulation in Colon Cancer
Huntington's Disease Signaling
EIF2 Signaling
Inhibition of ARE-Mediated mRNA Degradation Pathway
NRF2-mediated Oxidative Stress Response
Unfolded protein response
Molecular chaperones
Proteasome bounce-back
mRNA metabolism
Mitochondrial metabolism
Autophagy
Translation
Unfolded Protein Response
Oncogenic signal transducers
Cell motility
mRNA splicing
Protein Ubiquitination Pathway
FAT10 Signaling Pathway
BAG2 Signaling Pathway
Inhibition of ARE-Mediated mRNA Degradation Pathway
Huntington's Disease Signaling
Polyamine Regulation in Colon Cancer
Sirtuin Signaling Pathway
NRF2-mediated Oxidative Stress Response
Mitochondrial Dysfunction
Ferroptosis Signaling Pathway
Autophagy
Aldosterone Signaling in Epithelial Cells
Regulation of eIF4 and p70S6K Signaling
Cell Cycle Control of Chromosomal Replication
Unfolded protein response
Estrogen Receptor Signaling
EIF2 Signaling
tRNA Charging
14-3-3-mediated Signaling
Germ Cell-Sertoli Cell Junction Signaling
Cell Cycle: G2/M DNA Damage Checkpoint Regulation
Spliceosomal Cycle
Natural Killer Cell Signaling
Senescence Pathway
Hypoxia Signaling in the Cardiovascular System
Proteasome bounce-back
Molecular chaperones
Translation
Unfolded Protein Response
Apoptosis
Cell cycle progression
Unfolded protein response
Protein Ubiquitination Pathway
BAG2 Signaling Pathway
Huntington's Disease Signaling
EIF2 Signaling
Role of PKR in Interferon Induction and Antiviral Response
Endoplasmic Reticulum Stress Pathway
Death Receptor Signaling
Induction of Apoptosis by HIV1
Apoptosis Signaling
Kinetochore Metaphase Signaling Pathway
Cell Cycle: G2/M DNA Damage Checkpoint Regulation
for -log (B-H p-value<0.05)
Measurement: -log(B-H p-value) 0.00E00 9.3
B
Normalized to DMSO-CTRL
DMSO cancer
DMSO fibroblasts
Viability
CTRL
CHAPERONE MIX
HSPA1A/A1B
DNAJB1
HSP90AA1
HSPA8
AUTOPHAGY MIX
ATG5
CTSA
CTSD
ENDOCYTOSIS MIX
CHMP5
RAB11A
STAM3
CHAPERONES
AUTOPHAGY
ENDOCYTOSIS
Normalized to CFZ-CTRL
CFZ cancer
CFZ fibroblasts
C
CTRL
ATG5_1
ATG5_2
RAB11A_1
RAB11A_2
CHMP5_1
CHMP5_2
ATG5
RAB11A
CHMP5
GAPDH
CTRL
CTSD_1
CTSD_2
HSP90_1
HSP90_2
HSPA8
HSP90AA1
HSPA8
pro-CTSD
CTSD
GAPDH
CTRL
DNAJB1_1
DNAJB1_2
HSPA1A/B_1
HSPA1A/B_2
DNAJB1
HSPA1A/B
GAPDH

**Figure S2**

**A,** Ingenuity Pathway Analysis (IPA) of molecular pathways regulated by carfilzomib treatment, based on a differential analysis (carfilzomib vs. DMSO) of the proteomics data in Fig. 1G. Only proteins changing levels with a significance $p<0.05$ were used for analysis. Separate IPA core analyses were performed for each cell line and -log (B-H p-values) for canonical pathways were compared in IPA separately for cancer, normal fibroblasts and multiple myeloma cell lines, as indicated by captions, with a cutoff: B-H p-value<0.05. Colored captions indicate functional protein groups in the IPA pathways (indicated with arrows), selected for an siRNA vulnerability mini-screen in Fig. 2 B-C. Groups are marked in colors - specific to cancer (red) and specific multiple myeloma (magenta), pathways common to two of them (dark yellow, orange), or common to all studied cell types (light green).

**B,** Viability of the cancer cells and normal fibroblasts transfected with a mix or individual siRNAs targeting genes in the functional groups performing best in siRNA mini-screen – molecular chaperones, autophagy or endocytosis - treated with CFZ or DMSO for 48h. Mix of siRNAs contains all siRNAs used separately within a given functional group. Graphs present means of n=10, error bars represent SEM. Two-way ANOVA with Sidak correction for the cancer vs. fibroblasts comparison; $*p < 0.05$, $**p < 0.01$, $***p < 0.001$.

**C,** Western blot analysis of the silencing of the genes targeted in Fig. 2 D and Fig. S2 B

Figure S3

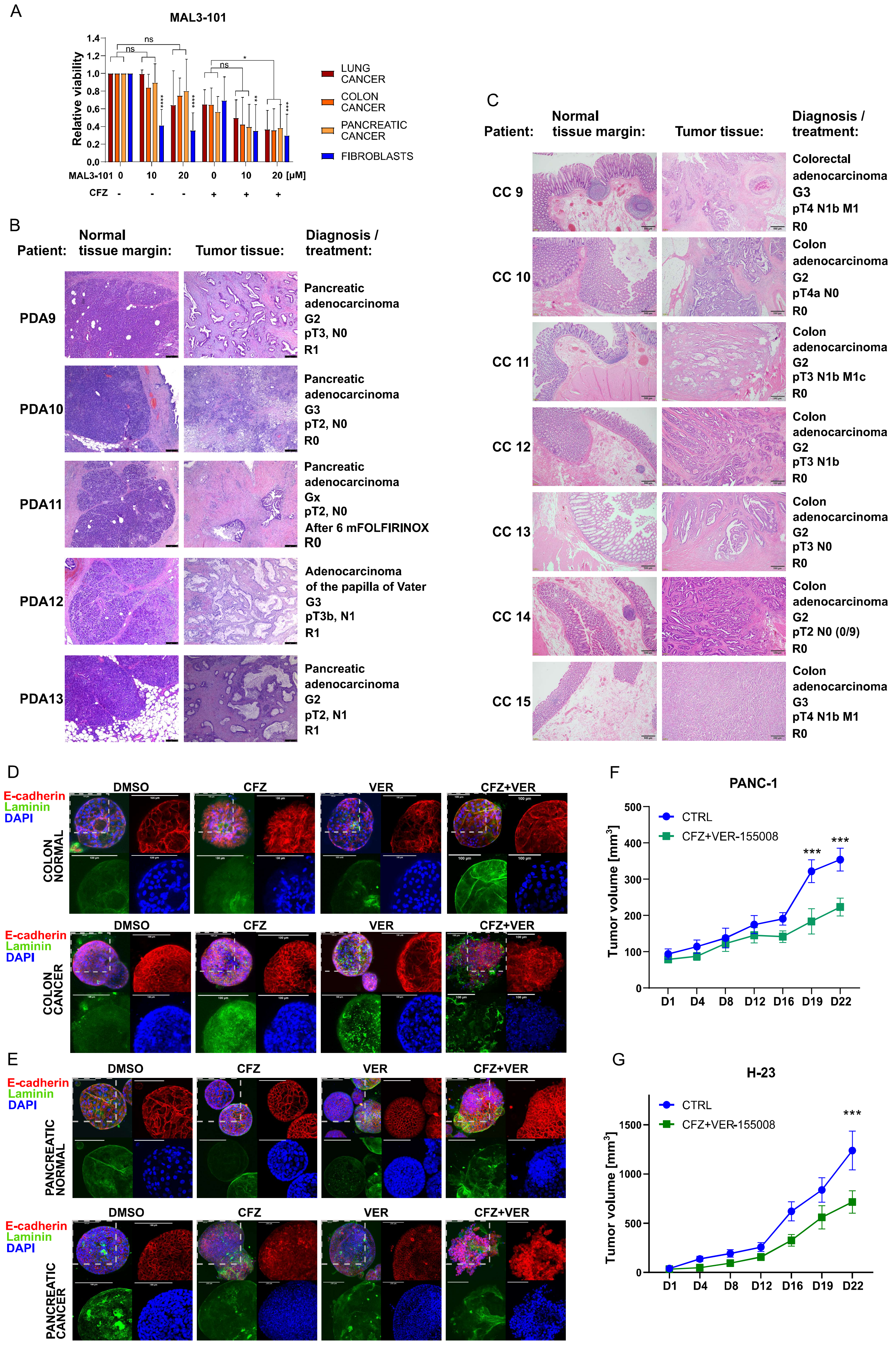
A
MAL3-101
Relative viability
1.4
1.2
1.0
0.8
0.6
0.4
0.2
0.0
ns
ns
ns
*
****
****
**
***
MAL3-101 0 10 20 0 10 20 [µM]
CFZ - - - + + +
LUNG CANCER
COLON CANCER
PANCREATIC CANCER
FIBROBLASTS
B
Patient:
Normal tissue margin:
Tumor tissue:
Diagnosis / treatment:
PDA9
Pancreatic adenocarcinoma
G2
pT3, N0
R1
PDA10
Pancreatic adenocarcinoma
G3
pT2, N0
R0
PDA11
Pancreatic adenocarcinoma
Gx
pT2, N0
After 6 mFOLFIRINOX
R0
PDA12
Adenocarcinoma of the papilla of Vater
G3
pT3b, N1
R1
PDA13
Pancreatic adenocarcinoma
G2
pT2, N1
R1
C
Patient:
Normal tissue margin:
Tumor tissue:
Diagnosis / treatment:
CC 9
Colorectal adenocarcinoma
G3
pT4 N1b M1
R0
CC 10
Colon adenocarcinoma
G2
pT4a N0
R0
CC 11
Colon adenocarcinoma
G2
pT3 N1b M1c
R0
CC 12
Colon adenocarcinoma
G2
pT3 N1b
R0
CC 13
Colon adenocarcinoma
G2
pT3 N0
R0
CC 14
Colon adenocarcinoma
G2
pT2 N0 (0/9)
R0
CC 15
Colon adenocarcinoma
G3
pT4 N1b M1
R0
D
E-cadherin
Laminin
DAPI
DMSO
CFZ
VER
CFZ+VER
COLON NORMAL
COLON CANCER
E
PANCREATIC NORMAL
PANCREATIC CANCER
F
PANC-1
Tumor volume [mm³]
500
400
300
200
100
0
CTRL
CFZ+VER-155008
***
***
D1 D4 D8 D12 D16 D19 D22
G
H-23
1500
1000
500
0
***
D1 D4 D8 D12 D16 D19 D22

**Figure S3**

**A,** Viability of cancer cell lines (H-23, H1299 – lung; PANC-1, CAPAN-2 – pancreatic; RKO, DLD-1 – colon) and normal fibroblasts treated with carfilzomib (CFZ), and HSPA1A/B inhibitor MAL3-101. Viability was measured with ATPlite after 48h of treatment with indicated drug concentrations. Error bars represent SEM. 2-way ANOVA with Tukey's correction, $*p < 0.05$; $**p < 0.01$; $***p < 0.001$;

**B, C,** Hematoxylin-Eosin (H&E) staining and diagnosis in pancreatic cancer (PDA) or colon cancer (CC) samples and patient-matched normal tissue margin samples used to generate organoid cultures in Fig. 3 B-E and Fig. S6 E. Histopathologic /clinical diagnosis and neoadjuvant treatment (used if indicated) for the cancer tissue is described in the diagnosis/treatment column.

**D, E,** Fluorescent microscopy of colon/pancreatic cancer and colon/pancreas normal organoids. Organoids were treated with carfilzomib and VER-155008 as indicated for 24h and fixed for staining. Red – E-cadherin; Green – Laminin-5; Blue – DAPI. Scale bar represents 100 µm. Grey dashed square marks a fragment of the merged picture which is enlarged in a single-channel pictures. Corresponding phase contrast microscopy pictures of pancreatic colon and normal organoids are included in supplementary figure (Fig. S3 C)

**F, G,** The size of xenografts formed in mice from PANC-1 and H-23 cancer cells. Indicated drugs were administered intraperitoneally**:** DMSO – control group, carfilzomib (CFZ) 4 mg/kg, VER-155008 35 mg/kg, VER+CFZ 35 mg/kg + 4 mg/kg, diluted in 200µl PBS-5% Tween-80 every other day. N = 7 mice / group in PANC-1 H-23 – 2x5 animals.

Figure S4

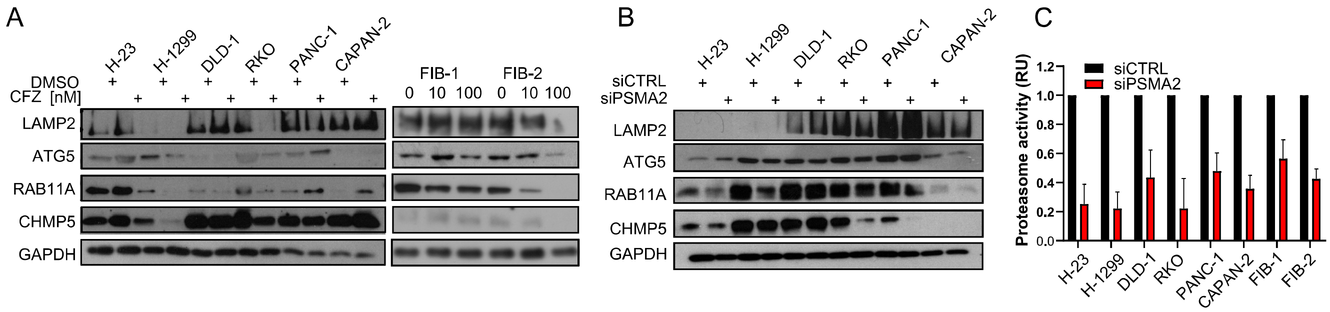


D

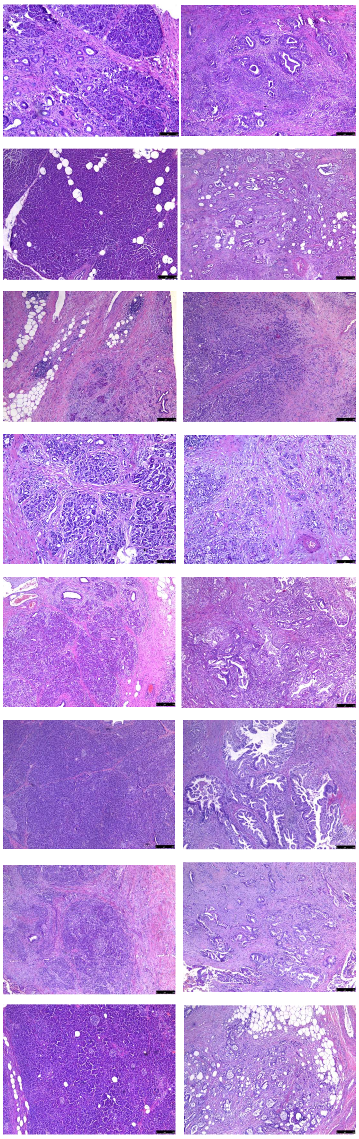

| Patient: | Normal tissue margin: | Tumor tissue: | Diagnosis / treatment: |
|---|---|---|---|
| PDA1 | | | Pancreatic adenocarcinoma G3 pT2, N1 R0 |
| PDA2 | | | Pancreatic adenocarcinoma G2 pT2, N2 R1 |
| PDA3 | | | Pancreatic adenocarcinoma G3 pT2, N0 R0 |
| PDA4 | | | Pancreatic adenocarcinoma G3 pT3, N1 R1 |
| PDA5 | | | Pancreatic adenocarcinoma G2 pT3, N2 R1 |
| PDA6 | | | Pancreatic adenocarcinoma G2 pT3, N1 R1 |
| PDA7 | | | Pancreatic adenocarcinoma G2 pT2, N1 R1 |
| PDA8 | | | Pancreatic adenocarcinoma G1 pT2, N0 R1 |

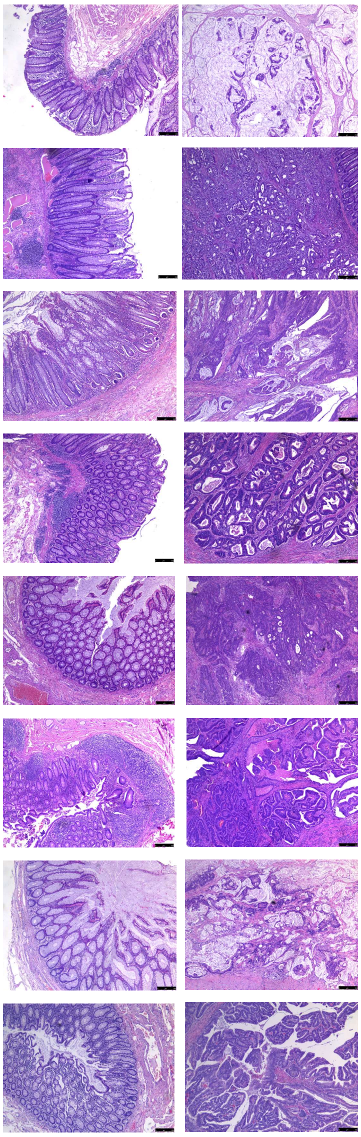

| Patient: | Normal tissue margin: | Tumor tissue: | Diagnosis / treatment: |
|---|---|---|---|
| CC 1 | | | Mucinous cecal adenocarcinoma pT4a, N2a, M1 |
| CC 2 | | | Colon adenocarcinoma G2 pT4a, N2b M1a |
| CC 3 | | | Colon adenocarcinoma G2 pT2, N1b R0 |
| CC 4 | | | Colon adenocarcinoma G2 pT3, N0 R0 |
| CC 5 | | | Adenocarcinoma of the rectosigmoid flexure G2 pT2, N1a R0 |
| CC 6 | | | Cecal adenocarcinoma G2 pT2, N1b R0 |
| CC 7 | | | Mucinous sigmoid adenocarcinoma G2 pT3, N0 R0 |
| CC 8 | | | Mucinous colon adenocarcinoma pT3, N0 R0 |

**Figure S4**

**A,** Autophagy and endocytosis related protein levels in cancer and normal fibroblast cell lines treated with carfilzomib for 24h (Concentrations as in Figure 1E unless stated otherwise) determined by western blot

**B,** Autophagy and endocytosis related protein levels in cancer cell lines transfected with siRNA targeting the proteasome subunit PSMA2, essential for the proteasome activity determined by western blot.

**C,** Bar graph presenting chymotrypsin-like proteasome activity on PSMA2 silencing. Bars are means of n=3 measurements with SD

**D,** Hematoxylin-Eosin (H&E) staining and diagnosis in pancreatic cancer (PDA) or colon cancer (CC) samples and patient-matched normal tissue margin samples used for protein and RNA extraction employed in Fig. 4 F-H and Fig. S4 D. Histopathologic /clinical diagnosis and neoadjuvant treatment (used if indicated) for the cancer tissue is described in the diagnosis/treatment column.

Figure S5

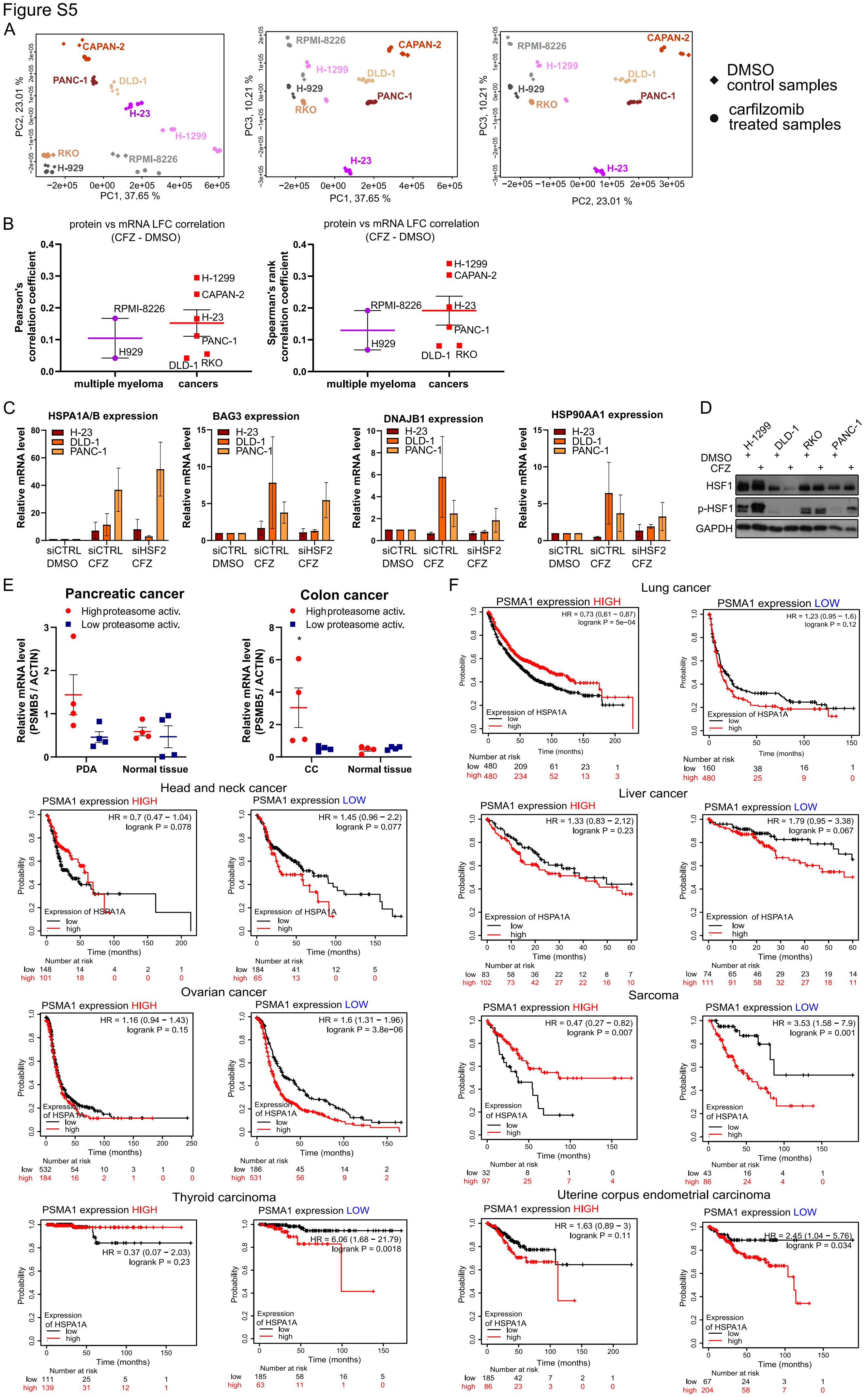

A
CAPAN-2
PANC-1
DLD-1
H-23
H-1299
RKO
H-929
RPMI-8226
PC1, 37.65 %
PC2, 23.01 %
PC3, 10.21 %
DMSO control samples
carfilzomib treated samples
B
protein vs mRNA LFC correlation (CFZ - DMSO)
Pearson's correlation coefficient
Spearman's rank correlation coefficient
multiple myeloma
cancers
C
HSPA1A/B expression
BAG3 expression
DNAJB1 expression
HSP90AA1 expression
Relative mRNA level
siCTRL DMSO
siCTRL CFZ
siHSF2 CFZ
D
H-1299
DLD-1
RKO
PANC-1
DMSO
CFZ
HSF1
p-HSF1
GAPDH
E
Pancreatic cancer
Colon cancer
High proteasome activ.
Low proteasome activ.
Relative mRNA level (PSMB5 / ACTIN)
PDA
Normal tissue
CC
F
Lung cancer
Head and neck cancer
Liver cancer
Ovarian cancer
Sarcoma
Thyroid carcinoma
Uterine corpus endometrial carcinoma
PSMA1 expression HIGH
PSMA1 expression LOW
Expression of HSPA1A
low
high
Probability
Time (months)
Number at risk

**Figure S5**

**A,** PCA (principal component analysis) in RNA-sequencing samples for cell lines indicated in the graphs, under control conditions (DMSO, D) or carfilzomib treatment (C).

**B,** Pearson and Sperman correlation coefficient scores of the log fold changes of quantified proteins (Fig. 1) *vs*. their coding transcripts (Fig. 5), plotted as means of indicated multiple myeloma cell lines (magenta) and cancer cell lines (red). Horizontal bold colored lines are means with SEM.

**C,** Expression of HSPA1A/B, BAG3, HSP90AA1, DNAJB1 mRNA in the presence of carfilzomib (CFZ) and silenced HSF2 (siHSF2). H-23, DLD-1 and PANC-1 cancer cells were transfected with siHSF2 or siCTRL (negative control) and afterwards treated with carfilzomib (concentrations as in Fig. 1E). Expression of indicated genes was analyzed by qPCR. Bars represent mean of n=3 with SD. Two-way ANOVA with Tukey's correction: $*p<0.05$; $**p<0.01$; $***p<0.001$.

**D,** Western blot analysis of HSF1 phosphorylation in the presence of carfilzomib in the indicated cancer cell lines.

**E,** Proteasome subunit PSMB5 mRNA relative levels in "high" and " low" proteasome activity sample groups from Fig. 5F, in cancer and normal margin tissues; PDA – pancreatic adenocarcinoma, CC – colorectal carcinoma;

**F,** Association of the HSPA1A/B expression with patient's survival in indicated cancer patients datasets - in samples with high or low (above/below median) expression of PSMA1, representing the proteasome expression. Red curve – high level of HSPA1A/B; Black curve - low level of HSPA1A/B. HR—hazard ratio; log-rank P—log-rank test P value for the curves comparison. Numbers below graphs indicate number of patients at risk (n) —total and at consecutive time points.

Figure S6

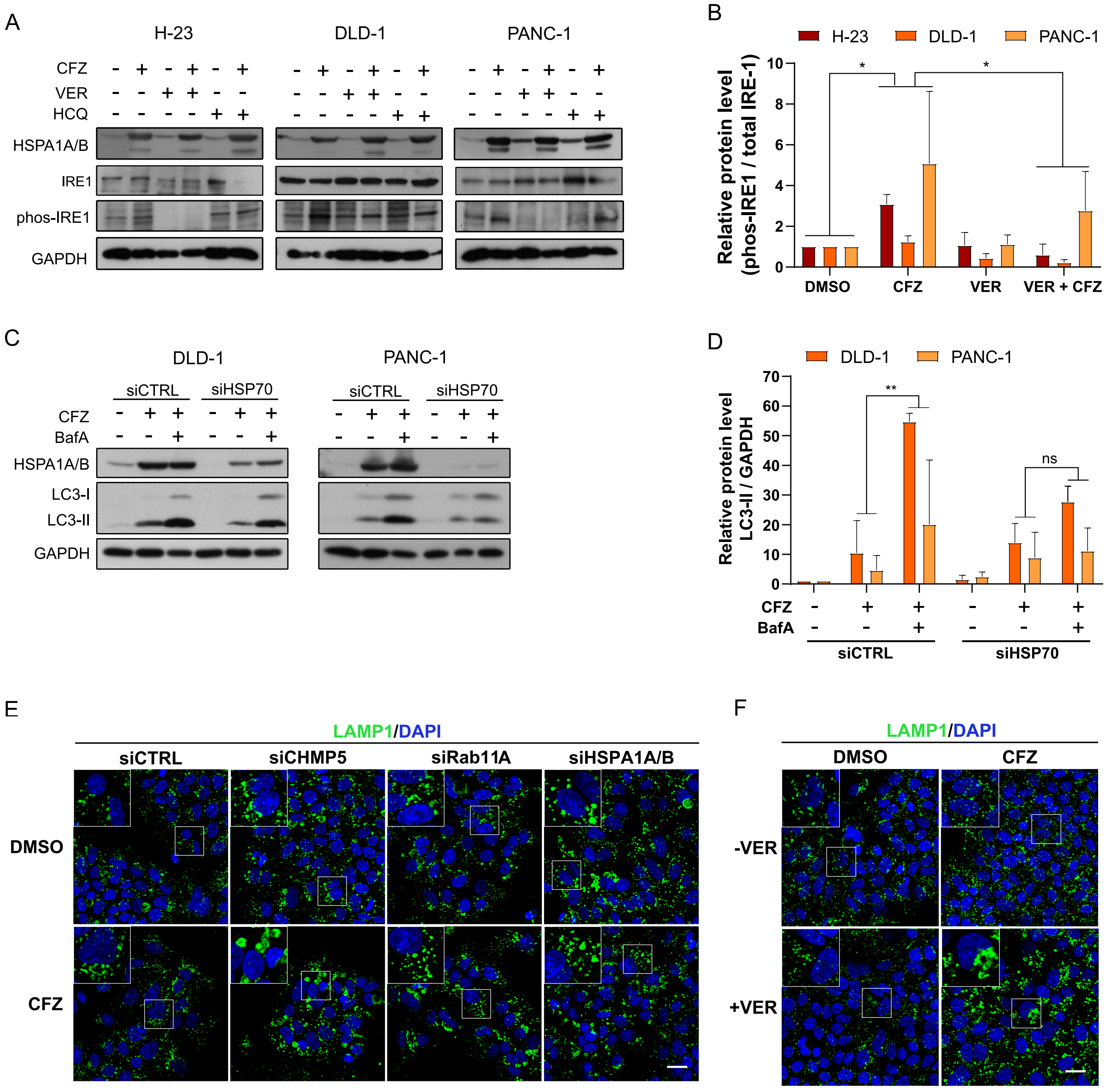

A
H-23
DLD-1
PANC-1
CFZ
VER
HCQ
HSPA1A/B
IRE1
phos-IRE1
GAPDH
B
H-23
DLD-1
PANC-1
Relative protein level (phos-IRE1 / total IRE-1)
DMSO
CFZ
VER
VER + CFZ
C
DLD-1
PANC-1
siCTRL
siHSP70
CFZ
BafA
HSPA1A/B
LC3-I
LC3-II
GAPDH
D
DLD-1
PANC-1
Relative protein level LC3-II / GAPDH
**
ns
CFZ
BafA
siCTRL
siHSP70
E
LAMP1/DAPI
siCTRL
siCHMP5
siRab11A
siHSPA1A/B
DMSO
CFZ
F
LAMP1/DAPI
DMSO
CFZ
-VER
+VER

**Figure S6**

**A,** IRE-1 phosphorylation on proteasome inhibition with carfilzomib and in combination with 50 µM VER-155008 or 80 µM hydroxychloroquine (HCQ). Indicated cells were treated for 24h. Carfilzomib concentrations as in Fig. 1F.

**B,** phospho-IRE-1 relative level on proteasome inhibition with carfilzomib and in combination with 40µM VER-155008 or 80µM hydroxychloroquine (HCQ). Bars represent mean of n=3 independent experiments, based on the western blot densitometry. Two-way ANOVA with Tukey's correction, $*p < 0.05$; $**p < 0.01$; $***p < 0.001$

**C,** LC3-I/LC3-II expression in indicated cell lines on HSPA1A/B (siHSP70) silencing and subsequent treatment with carfilzomib 100 nM (CFZ) bafilomycin A 20nM or control DMSO.

**D,** Densitometry-based LC3-II relative level on HSPA1A/B (siHSP70) silencing and subsequent treatment with carfilzomib 100 nM (CFZ), bafilomycin A 20 nM or control DMSO. Bars represent mean of n=2 with SD. Two-way ANOVA with Tukey's correction, ns $p > 0.05$; $**p < 0.01$;

**E, F,** Immunofluorescence staining of LAMP1 in DLD-1 colon cancer cell line transfected with indicated siRNAs and treated with 50nM CFZ (E) or treated with 50nM CFZ and 50µM VER-155008 (F) for 24h. Insets show magnified views of the regions boxed in the main images. Scale bars: 20 µm

Figure S7

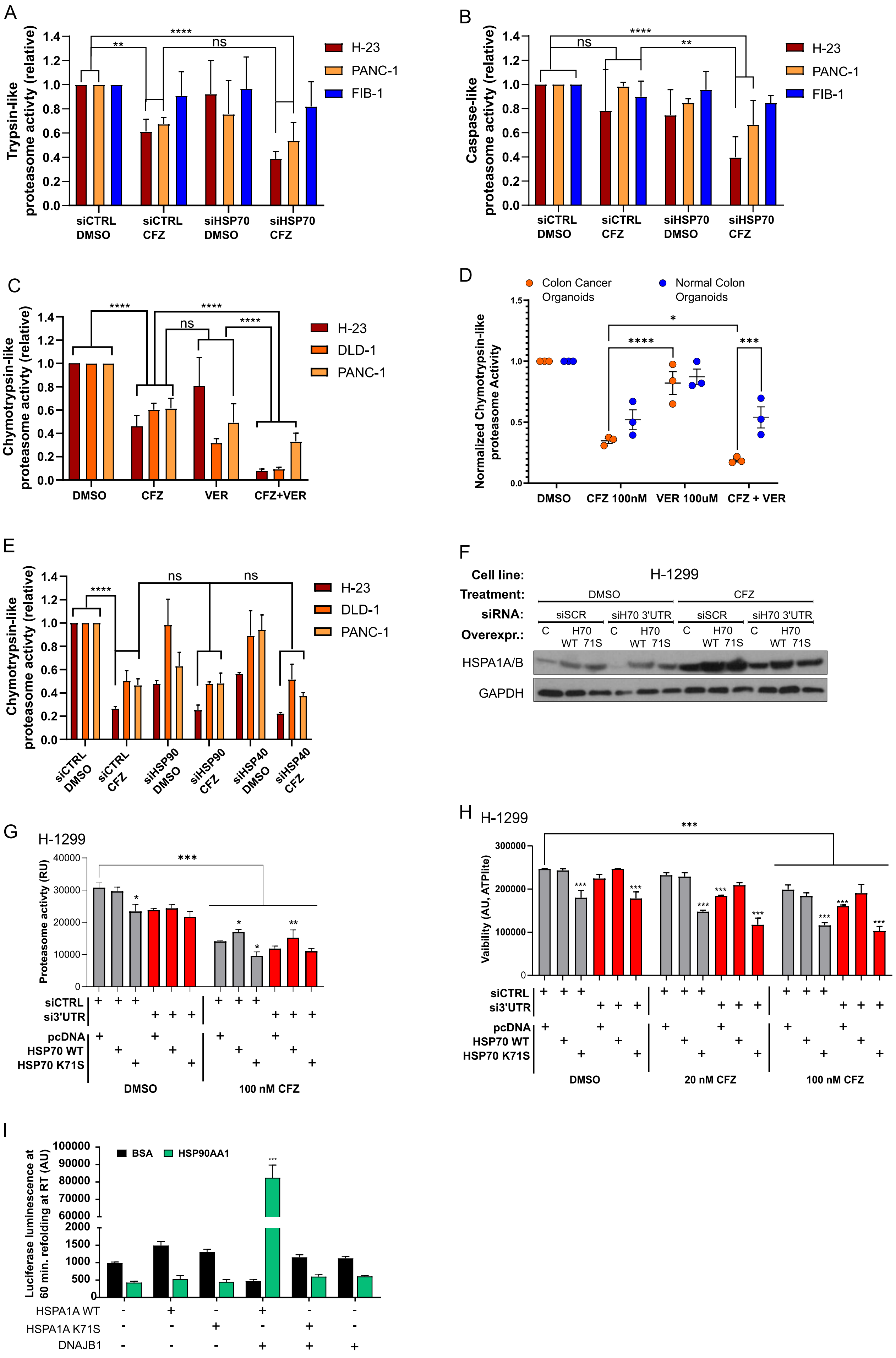
A
Trypsin-like proteasome activty (relative)
H-23
PANC-1
FIB-1
siCTRL DMSO
siCTRL CFZ
siHSP70 DMSO
siHSP70 CFZ
B
Caspase-like proteasome activty (relative)
H-23
PANC-1
FIB-1
C
Chymotrypsin-like proteasome activty (relative)
H-23
DLD-1
PANC-1
DMSO
CFZ
VER
CFZ+VER
D
Colon Cancer Organoids
Normal Colon Organoids
Normalized Chymotrypsin-like proteasome Activity
DMSO
CFZ 100nM
VER 100uM
CFZ + VER
E
Chymotrypsin-like proteasome activty (relative)
siCTRL DMSO
siCTRL CFZ
siHSP90 DMSO
siHSP90 CFZ
siHSP40 DMSO
siHSP40 CFZ
F
Cell line:
H-1299
Treatment:
DMSO
CFZ
siRNA:
siSCR
siH70 3'UTR
Overexpr.:
C
H70 WT 71S
HSPA1A/B
GAPDH
G
H-1299
Proteasome activty (RU)
siCTRL
si3'UTR
pcDNA
HSP70 WT
HSP70 K71S
DMSO
100 nM CFZ
H
H-1299
Vaibility (AU, ATPlite)
siCTRL
si3'UTR
pcDNA
HSP70 WT
HSP70 K71S
DMSO
20 nM CFZ
100 nM CFZ
I
Luciferase luminescence at 60 min. refolding at RT (AU)
BSA
HSP90AA1
HSPA1A WT
HSPA1A K71S
DNAJB1

**Figure S7**

**A,** Trypsin-like proteasome activity upon carfilzomib and silencing of HSPA1A/B (siHSP70) in the lung cancer (H-23), pancreatic cancer (PANC-1) and normal fibroblast (FIB-1) cell lines; siCTRL – negative siRNA control.

**B,** Caspase-like proteasome activity upon carfilzomib and silencing of HSPA1A/B (siHSP70) in the lung cancer (H-23), pancreatic cancer (PANC-1) and normal fibroblast (FIB-1) cell lines; siCTRL – negative control.

**C,** Chymotrypsin-like proteasome activity in the presence of carfilzomib and HSPA1A/B inhibitor VER-155008 in the lung cancer (H-23), colon cancer (DLD-1) and pancreatic (PANC-1) cell lines.

**D,** Chymotrypsin-like proteasome activity in the presence of carfilzomib and HSPA1A/B inhibitor VER-155008 in the 3 pairs of patient-matched colon cancer and normal tissue-derived organoid cultures.

**E,** Chymotrypsin-like proteasome activity upon carfilzomib and silencing of HSP90AA1 (siHSP90) or DNAJB1 (siHSP40) in the lung cancer (H-23), colon cancer (DLD-1). pancreatic cancer (PANC-1) cell lines; siCTRL – negative control.

**F,** Representative western blot demonstrating the silencing and the rescue overexpression of *HSPA1A/B* in H-1299 lung cancer cell line. siSCR – negative control; siH70 3'UTR – siRNA targeting 3'UTR of *HSPA1A/B* mRNA; H70 WT and H70 71S – vectors expressing HA-tagged wild type and K71S mutant of HSPA1A, respectively. Treatment with carfilzomib (CFZ) using concentrations as in Fig. 1E.

**G,** Chymotrypsin-like proteasome activity in H-1299 lung cancer cells upon silencing of *HSPA1A/B* and rescue overexpression of wild type (WT) or mutant (K71S) HSPA1A in the presence of indicated carfilzomib concentration. Bars are means of n=2 with SD.

**H,** Viability of H-1299 lung cancer cells upon silencing of HSPA1A/B and rescue overexpression of wild type (WT) or mutant (K71S) HSPA1A in the presence of indicated carfilzomib concentrations. Bars are means of n=3 with SD.

**I,** Refolding of heat inactivated, purified firefly luciferase in the presence of indicated chaperone proteins from HSP70, HSP40 and HSP90 families (for details see Methods). Bars are means of n=3 with SD. Two-way ANOVA with Tukey's correction, $*p < 0.05$; $**p < 0.01$; $***p < 0.001$

### Figure S8

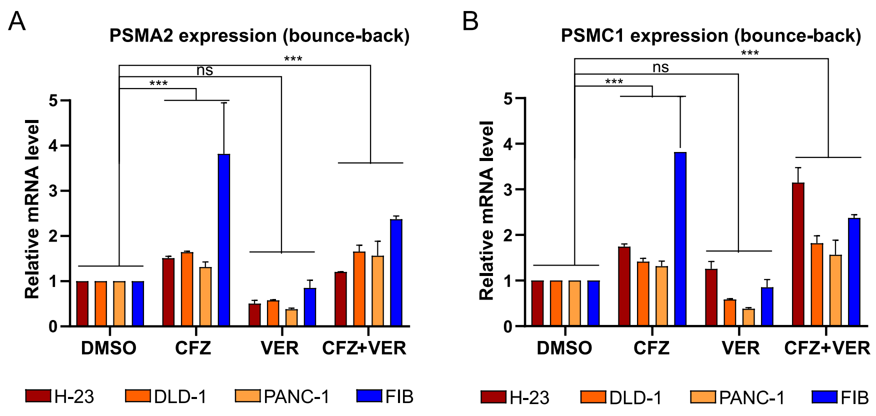
A
PSMA2 expression (bounce-back)
Relative mRNA level
0
1
2
3
4
5
***
ns
***
DMSO
CFZ
VER
CFZ+VER
H-23
DLD-1
PANC-1
FIB
B
PSMC1 expression (bounce-back)
Relative mRNA level
0
1
2
3
4
5
***
ns
***
DMSO
CFZ
VER
CFZ+VER
H-23
DLD-1
PANC-1
FIB

**Figure S8**

**A, B,** Proteasome subunits *PSMA2* and *PSMC1* transcriptional bounce-back upon carfilzomib and HSPA1A/B inhibition with VER-155008 in the indicated lung (H-23), colon (DLD-1) and pancreatic (PANC-1) cancer and normal fibroblast cell lines; siSCR – negative control. Bars are means of n=2 with SD.
